# Paired primary-metastasis patient-derived organoids and mouse models identify phenotypic evolution and druggable dependencies of peritoneal metastasis from appendiceal cancer

**DOI:** 10.1101/2025.02.17.638725

**Authors:** Ahmed Mahmoud, Philip H. Choi, Christine Sukhwa, Jura Pintar, Henry Walch, Nan Zhao, Jonathan Bermeo, Sebastian Chung, Manisha Raghavan, Samhita Bapat, Qingwen Jiang, Georgios Karagkounis, Julia Meredith, Michael Giarrizzo, Canan Firat, Andrea Cercek, Michael B. Foote, Nikolaus Schultz, Walid K. Chatila, Garrett M. Nash, Jinru Shia, Francisco Sanchez-Vega, Steven Larson, Arvin C. Dar, Neal Rosen, Karuna Ganesh

## Abstract

Peritoneal carcinomatosis is a common yet deadly manifestation of gastrointestinal cancers, with few effective treatments. To identify targetable determinants of peritoneal metastasis, we focused on appendiceal adenocarcinoma (AC), a gastrointestinal cancer that metastasizes almost exclusively to the peritoneum. Current treatments are extrapolated from colorectal cancer (CRC), yet AC has distinct genomic alterations, mucinous morphology and peritoneum restricted metastatic pattern. Further, no stable preclinical models of AC exist, limiting drug discovery and representing an unmet clinical need. We establish a first-in-class stable biobank of 16 long-term cultured AC patient-derived organoids (PDOs), including 3 matched, simultaneously resected primary AC-peritoneal carcinomatosis (AC-PC) pairs. By enriching for cancer cells, AC PDOs enable accurate genomic characterization relative to paucicellular AC tissue. We establish an organoid orthotopic intraperitoneal xenograft model that recapitulates diffuse peritoneal carcinomatosis and show that PC-organoids retain increased metastatic capacity, decreased growth factor dependency and sensitivity to standard of care chemotherapy relative to matched primary AC organoids. Single cell profiling of AC-PC pairs reveals dedifferentiation from mucinous differentiated states in primary AC into intestinal stem cell and fetal progenitor states in AC-PC, with upregulation of oncogenic signaling pathways. Through hypothesis-driven drug testing, we identify KRAS^MULTI^-ON inhibitor RMC-7977 and Wnt-targeting tyrosine kinase inhibitor WNTinib as novel, clinically actionable strategies to target AC-PC more effectively.

## Introduction

The peritoneal cavity is a common site of metastasis from advanced gastrointestinal and gynecological malignancies, up to 70% of patients present with peritoneal carcinomatosis at the time of metastatic diagnosis^1–3^. Peritoneal carcinomatosis (PC) can lead to the loss of intestinal peristalsis and organ failure, and poor overall survival of just 3-24 months from diagnosis^4–6^. Further, peritoneal metastasis demonstrates poor response to standard-of-care systemic chemotherapy in comparison to extraperitoneal metastasis^7,8^. Cytoreductive surgery and intraperitoneal chemotherapy have thus been used to directly target peritoneal metastasis in patients with low tumor burden, yet, in many patients, they typically only accomplish temporary disease control and can be associated with a high rate of often severe complications^9,10^. For patients with widespread carcinomatosis, few effective therapeutic strategies exist. Progress in developing effective treatment strategies has been limited by the lack of clinically representative preclinical models that accurately capture the heterogenous tumor cell states, frequently mucinous histology, and diffuse distribution of peritoneal metastases seen in patients, where tumor deposits can coat much of the intra-abdominal structures and serosal surfaces.

Despite the lack of prospective clinical trials demonstrating a survival benefit from this approach, PC patients undergo extensive cytoreductive surgery and, frequently, intraperitoneal chemotherapy, but have a median survival of less than 48 months^11,12^. Many gastrointestinal cancers metastasize concurrently through the lymphovascular and hematogenous routes to distant organs and through transcoelomic spread directly to the peritoneum; therefore, studies on PC are often confounded by the presence of non-PC metatstases^13,14^. Appendiceal adenocarcinoma (AC) is an archetypal gastrointestinal cancer that metastasizes largely to the peritoneum^15^. AC is frequently diagnosed incidentally during appendectomies, which may be curative for primary AC, but >25% of patients present with metastatic disease at the time of diagnosis^16^. AC-PC is thought to arise from transcoelomic spread as cancer cells invade through the appendiceal wall and colonize adjacent peritoneum ^17–19^. AC patients are managed using systemic chemotherapies that have been shown to be effective in patients with colorectal cancer (CRC) metastasis to the liver and lungs, but these regimens have shown limited efficacy in PC from both AC and CRC^20–22^. Moreover, AC tumors exhibit distinct pathology, histology, and mutational profile from CRC^23–26:^ AC exhibits fewer APC mutations (5% vs 79%) and more frequent KRAS (58% vs 44%) and GNAS (33% vs 5%) mutations than CRC^23,27,28^. These data underscore the need for representative preclinical models to identify effective therapeutic strategies for this underserved patient population. One previous study treated short-term 7-day cultures of AC and immune cells from 19 patients with immune checkpoint blockade^29^. Another study grew organotypic slice cultures from 6 AC patients for 7 days and demonstrated heterogenous response to immunomodulator TAK981^30^. One recent study established three AC patient-derived xenografts (PDX) from 16 patients^31^, but there remains a lack of well-characterized, stable cultures of primary AC or AC-PC capable of expansion and long-term passaging, to enable scalable preclinical drug testing and mechanistic interrogation, representing a critical unmet need.

To address these challenges, we generated and characterized a stable long-term AC patient-derived organoid biobank and a novel organoid orthotopic intraperitoneal xenograft model of diffuse peritoneal carcinomatosis. Developing a unique resource of paired primary AC and AC-PC PDOs derived from simultaneously surgically resected AC primary tumor-metastasis tissue from the same patients, we demonstrate that PC-organoids retain increased metastatic capacity and dedifferentiated, stem-like transcriptional states, decreased exogenous growth factor dependency and sensitivity to standard of care chemotherapy relative to matched primary AC organoids. Through hypothesis-driven drug testing, we identify the RAS inhibitor RMC-7977 and Wnt-targeting tyrosine kinase inhibitor WNTinib as novel, clinically actionable strategies that significantly reduce AC-peritoneal carcinomatosis *in vivo*.

## Results

### An appendiceal cancer organoid biobank recapitulates patient tumor morphology and histopathology

We collected surgically resected primary and metastatic tumors from 24 patients with AC undergoing appendectomies and/or cytoreductive surgeries at Memorial Sloan Kettering Cancer Center. Metastatic tissue was collected from the parietal peritoneum, omentum and ovaries. Tumors were dissociated into single cell suspension and plated in Matrigel in human intestinal stem cell (HISC) media to generate organoid cultures (n = 23) or mixed with 50% Matrigel and implanted subcutaneously into immunocompromised mice to generate patient-derived xenografts (n=3) (**Figure 1a**). 14/23 initial organoid cultures could be successfully passaged >5 times, cryopreserved and successfully thawed and regrown as stable long-term cultures. 9 additional tumors yielded short term cultures that did not survive freeze-thaw. 2/3 xenografts were harvested, dissociated and could be subsequently grown and passaged as long-term organoid culture. A total of 16 long-term patient-derived organoid cultures were established, including three primary-metastasis matched pairs established synchronously from the same surgery (**Figure 1a, b**). We had the greatest success establishing organoids from poorly differentiated tumors (14/15), while cells from moderately-(1/10) and well-differentiated (1/5) tumors as well as normal appendix largely failed to generate stable organoids in HISC media (**Table S1**).

**Figure 1.**
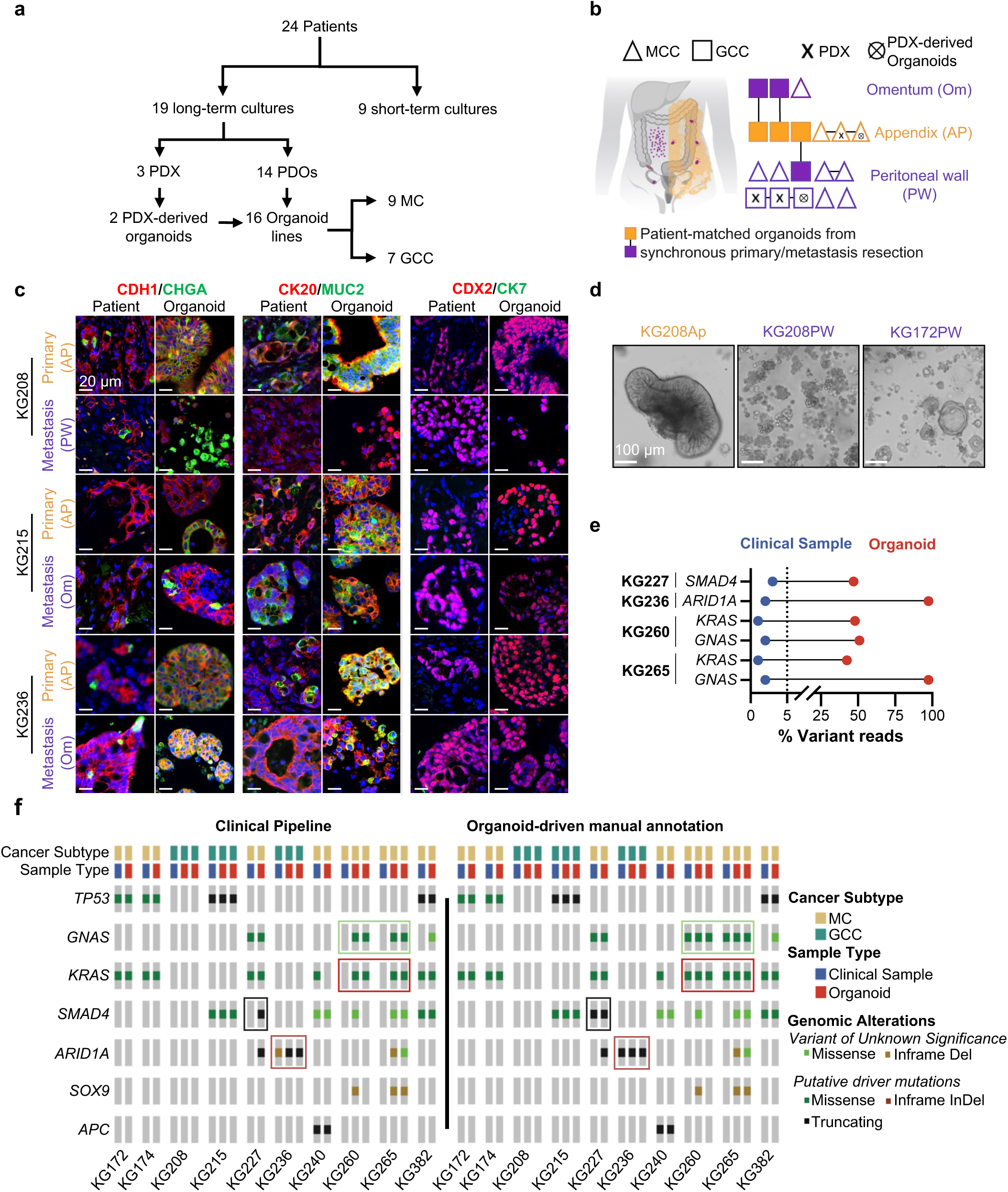
**Establishment of a long-term, histopathologically and genomically representative appendiceal cancer organoid biobank. a,b**, Schematic diagram of appendiceal cancer biobank. PDX: patient-derived xenograft; PDO: patient-derived organoids; MC: mucinous carcinoma; GCC: goblet cell carcinoma. **b**, Lines represent patient-matched samples. Shaded boxes represent patient-matched synchronously resected primary:metastasis PDOs. **c**, Representative immunofluorescence staining of paired primary tumor (yellow label) and metastasis (purple label)-derived GC AC organoids derived from three patients, and corresponding parental tumors, showing concordant cell types conserved across tumor and organoid. Left: CDH1 (red) & CHGA (green); Center: KRT20 (red) & MUC2 (green); Right: CDX2 (red) & CK7 (green). Nuclei are counterstained with DAPI (blue). **d**, Representative brightfield images of primary and metastasis organoids. **e,** Percent variant reads of mutations found through manual annotations in organoids and clinical samples. Dotted line represents the cutoff for the MSK-IMPACT clinical pipeline. **f**, Mutational landscape of 23 AC tumors and matched organoids, detected by MSK-IMPACT targeted exon sequencing. Tumor samples were processed through the standard pipeline (left) or reannotated by manually examining aligned reads based on organoid mutation calls (right) to call mutations in paucicellular clinical AC samples.

Epithelial AC tumors are histopathologically broadly classified as mucinous (MC) or goblet cell (GCC) carcinoma, although such classification is purely morphology based and does not correlate well with genetics or clinical outcomes^32,33^. MCs appear paucicellular with a large extracellular mucinous component, resembling ‘low-grade’ well-differentiated tumors. GCCs consist of an epithelial component, similar to mucinous tumors, and a ‘neuroendocrine’ component which consists of cells exhibiting a signet-cell phenotype with a large cytoplasmic mucus vacuole which squeezes the nucleus to the side^32,34,35^. Pathology expert review of paired tumor and organoid hematoxylin and eosin (H&E) slides validated that the organoids recapitulate the morphological classification of their parental tumors, retaining the signet-cell phenotype found in many of the tumors (**Figure S1a**). Copious mucin production is a characteristic clinical feature of AC, leading to highly symptomatic accumulation of paucicellular dense ascites termed “pseudomyxoma peritonei”, and limiting drug penetration to mucin-protected tumor nests^36–39^. Alcian blue staining of matched tumors and organoids confirmed that AC organoids retain mucin production capacity (**Figure S1b**). Next, we asked whether AC organoids retain the immunophenotypic similarities and differences of AC tumors with respect to other common tumors that metastasize to the peritoneum, i.e. colorectal cancer (CRC) and ovarian cancer (OV). Similar to CRC tumors, both GCC (**Figure 1c**) and MC (**Figure S1c**) were positive for expression of the intestinal lineage-defining transcription factor CDX2 and epithelial differentiation markers CDH1 and CK20 (**Figure S1d**). CK7, expressed widely in ovarian cancer and in a subset of cells in CRC lung metastasis, was not expressed by any of the AC primary or metastatic tissues or organoids. Both MC and GCC AC express intracellular mucin (MUC2) and the neuroendocrine marker chromogranin A (CHGA). AC organoids exhibited diverse morphologies, from dense structures to mucin-filled lumina and branching budding-like structures (**Figure 1d**).

### AC organoids identify previously undetected mutations in paucicellular tumors

To determine whether the organoids recapitulate the genomic alterations of the corresponding patient tissue, we performed MSK Integrated Mutation Profiling of Actionable Cancer Targets (MSK-IMPACT)^40–42^. We surprisingly identified several mutations in the organoids that were not called in the patient samples using the standard clinical pipeline, including potentially clinically actionable *KRAS* mutations associated with poor prognosis^23^ (**Figure 1f**, left panel). Manual in-depth reannotation of the clinical MSK-IMPACT results revealed the corresponding presence of these mutations (**Figure 1e, 1f**, right panel). MSK-IMPACT utilizes a 5% variant allele cutoff to determine the presence of mutations. Since AC tumors are frequently paucicellular, with the bulk of the tumor mass comprised by mucin (**Figure S2a**), low tumor purity can decrease the sensitivity of clinical mutation testing, and therapeutically actionable mutations can be overlooked. These data highlight the utility of AC organoids as a means of enriching for cancer cells as a source of pure tumor DNA and a valuable tool for informing clinical decision-making and treatment selection.

### An organoid orthotopic model of diffuse peritoneal carcinomatosis reveals differential metastatic capacity of paired primary AC and AC-PC organoids

We next sought to develop a mouse model which recapitulates the slow and diffuse intracoelomic spread of peritoneal carcinomatosis. Upon intraperitoneal injection, most cancer cells typically form single focal solid tumors within the peritoneum, unlike PC in patients which presents as diffuse seeding throughout the peritoneum^15^. By titrating down the amount of Matrigel admixed with a CRC organoid cell suspension during intraperitoneal injection, followed by abdominal massage (see **Methods**), we established a clinically representative model of slow and diffuse peritoneal carcinomatosis (DPC), while not altering the overall tumor burden (**Figure S3a-c**).

Using our DPC model, we asked whether paired primary AC/AC-PC organoids established synchronously from the same patients retained distinct abilities to establish peritoneal metastasis. Upon transplantation of td-Tomato-AkaLuc labeled paired primary-metastasis paired organoids from three patients, all three metastasis organoids (KG208PW, KG215Om and KG236Om) successfully established DPC *in vivo*, while the corresponding matched primary tumor-derived organoids (KG208Ap, KG215Ap and KG236Ap) failed to colonize the peritoneum (**Figure 2a-c**). AC PDOs showed a marked preference for fatty tissues, growing within the omentum as well as other adipocyte-rich sites including perirectal and retroperitoneal fat (**Figure S3d**), consistent with the peri-adipocyte location of AC peritoneal deposits observed in the corresponding patient tissue (**Figure S3e**). DPC xenograft tumors recapitulated the signet-cell morphology, and the expression of epithelial, mucin and neuroendocrine markers observed in the corresponding patient tumors (**Figures 2d; S3e,f**).

**Figure 2.**
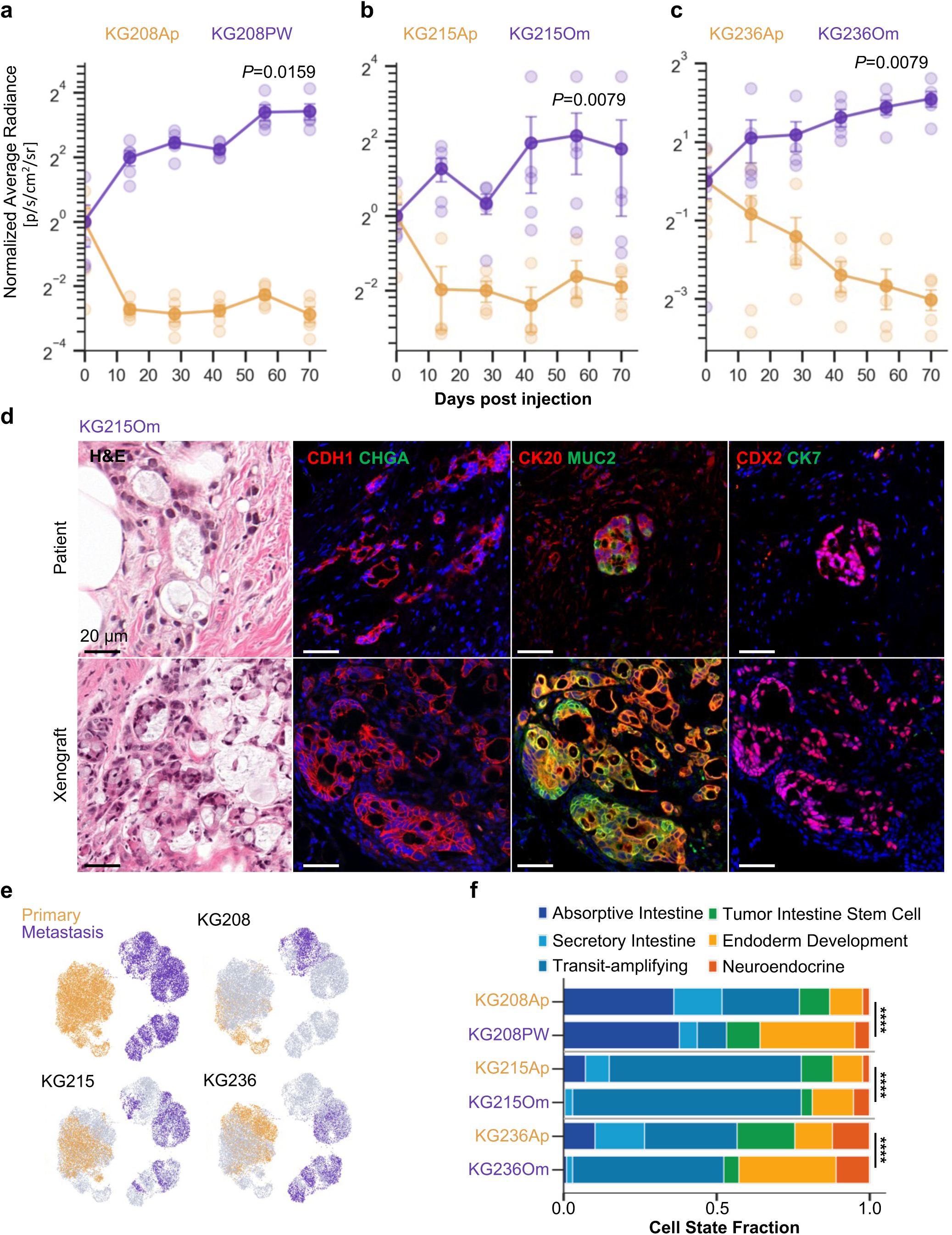
**An organoid orthotopic mouse model of diffuse peritoneal carcinomatosis demonstrates increased peritoneal metastasis capacity of PC organoids compared to paired primary AC organoids. a-c**, Normalized average radiance measured by biweekly *ex vivo* bioluminescence imaging following intraperitoneal injection of 500,000 cells from matched primary tumor (yellow) and peritoneal metastasis (purple) organoids from (**a**) KG208Ap and KG208PW, (**b**) KG215Ap and KG215Om, (**c**) KG236Ap and KG236Om lines in NSG mice, normalized to the mean average radiance across each group immediately following injection (day 0). Mean ± s.e.m. n = 4 (KG215Ap), 5 (KG215Om), 5 (KG208Ap), 5 (KG208PW), 4 (KG236Ap), 5 (KG236Om). Two-sided Mann Whitney rank-sum test. **d**, H&E and immunofluorescence staining with the indicated antibodies, of KG215Om diffuse peritoneal xenograft tumors and corresponding patient peritoneal metastasis tissue. **e,** scANVI latent space uniform manifold approximation and projection (scANVI-UMAP) plot of the of KG208Ap, KG208PW, KG215Ap, KG236Ap KG215Om and KG236Om organoids (25,103 cells). **f**, The distribution of module fractions for KG208Ap/PW, KG215Ap/Om and KG236Ap/Om, labels are colored by sample’s origin, primary or metastasis. Significant differences are indicated by asterisks as calculated by two-sided chi-squared test **** p < 0.0001.

The retention of differential metastatic capacity by primary tumor and metastasis-derived AC organoids from the same patients suggests the enrichment of distinct cell-autonomous, epigenetically encoded phenotypic states during peritoneal metastasis. To delineate the cell states that confer peritoneal metastatic capacity in AC, we performed 10X single cell mRNA sequencing on three pairs of primary AC and AC-PC organoids (KG208Ap/PW, KG215Ap/Om, KG236Ap/Om) (**Figure 2e**). We recently discovered that the transition from primary tumor to liver metastasis involves a progressive loss of expression of canonical intestinal genes, accompanied by dedifferentiation into a fetal-like endoderm progenitor state and gain of expression of non-canonical neuroendocrine and squamous genes ^43^. We, therefore, asked whether a similar transition might be occurring during AC peritoneal metastasis. GSEA of pseudobulked AC primary-metastasis pairs showed loss of intestine-specific gene expression in metastasis (**Figure S4a**). Pairwise comparisons of primary and metastasis organoids showed a significant shift of cell state composition, with metastasis organoids containing a smaller fraction of differentiated intestinal cells and a higher fraction of cells with endoderm development and neuroendocrine gene expression (**Figure 2f, S4a, b**). Thus, as in CRC liver metastasis, AC peritoneal metastasis selects against canonical intestinal gene expression, and instead selects for expression of fetal endoderm-like and non-canonical neuroendocrine genes. Together our data suggest that shared programs of dedifferentiation and plasticity drive metastasis across diverse cancer cells of origin across the intestine and across the liver and peritoneum as metastatic sites.

### Increased niche independence of AC-PC organoids in comparison to paired primary AC organoids

Sustained cell-autonomous mitogenic signaling is a hallmark of cancer and is coupled with metastatic progression^44–46^. We therefore hypothesized that peritoneal metastasis might select for cancer cells with increased cancer cell-intrinsic mitogenic signaling pathway activation, that are capable of niche-independent growth. Indeed, in comparison to primary AC organoids, the expression of Wnt, PI3K/AKT and MYC target genes was significantly upregulated in AC-PC organoids from the same patients grown in identical *ex vivo* human intestinal stem media (containing Wnt, R-spondin, EGF, noggin, A83-01, FGF and IGF1)^47^ (**Figure S4c**). Next, we asked whether the increased intrinsic activation of mitogenic signaling pathways in AC-PC organoids would confer decreased sensitivity to extrinsic mitogen deprivation, enabling niche-independent growth in the mitogen-low peritoneal microenvironment. We sequentially withdrew from the organoid media individual growth factors or combinations known to activate key signaling pathways required for intestinal stem cell renewal and proliferation: Wnt/R-spondin to induce Wnt signaling, EGF to induce MAPK signaling, noggin/A83-01: to inhibit TGFβ signaling, FGF/IGF1 to induce PI3K/AKT and MEK/ERK signaling (**Figure 3a-c, S5**). Wnt pathway dysregulation, frequently through early genetic inactivation of APC, is a key step in CRC initiation as well as in hepatocellular carcinoma but is uncommon in AC^48–51^. Upon withdrawal of Wnt and R-spondin from growth media (ENAFI), both primary and metastasis AC organoids failed to proliferate, in contrast to APC-mutant CRC primary (CRC146P) and liver metastasis (CRC146Li), which were unaffected by exogenous Wnt deprivation (**Figure 3a, S5**). AC and CRC PDOs were unable to survive simultaneous depletion of three receptor tyrosine kinases (RTK) agonists, IGF1, FGF and EGF, the depletion of FGF and IGF1 or the depletion of FGF alone (**Figure 3a**). Primary AC organoid lines KG208Ap and KG236Ap showed reduced growth upon withdrawal of TGFβ inhibitors noggin and A83-01, but their metastatic AC-PC counterparts remained unaffected despite a lack of SMAD4 or other TGFβ pathways mutations assayed by MSK-IMPACT, similar to the SMAD4 mutant KG215 and CRC146 lines (**Figure 3a**). While we cannot rule out the acquisition of mutations not assayed by MSK-IMPACT, these data suggest that PC selects for functional insensitivity to TGFβ, acquired through either genetic or non-genetic mechanisms. The niche factors appeared to have differential effects on organoid initiation capacity of single cells (stemness) and organoid size (proliferation). Wnt/R-spondin withdrawal (ENAFI) resulted in both fewer and smaller AC organoids, suggesting a requirement for Wnt signaling for both stemness and proliferation in AC (**Figure 3b, d; S5a, b**). In contrast, CRC146P formed fewer but larger organoids in the absence of exogenous Wnt ligands (**Figure S5a, b**). On the other hand, EGF withdrawal led to the formation of more organoids (increased stemness) which were smaller (decreased proliferation), while FGF withdrawal decreased both stemness and proliferation (**Figure 3b, c; S5a, b**). Together, these results suggest that AC-PC selects for cancer cells with increased cell intrinsic mitogenic signaling that are less dependent on exogenous mitogenic factors for survival, metastatic seeding, and outgrowth.

**Figure 3.**
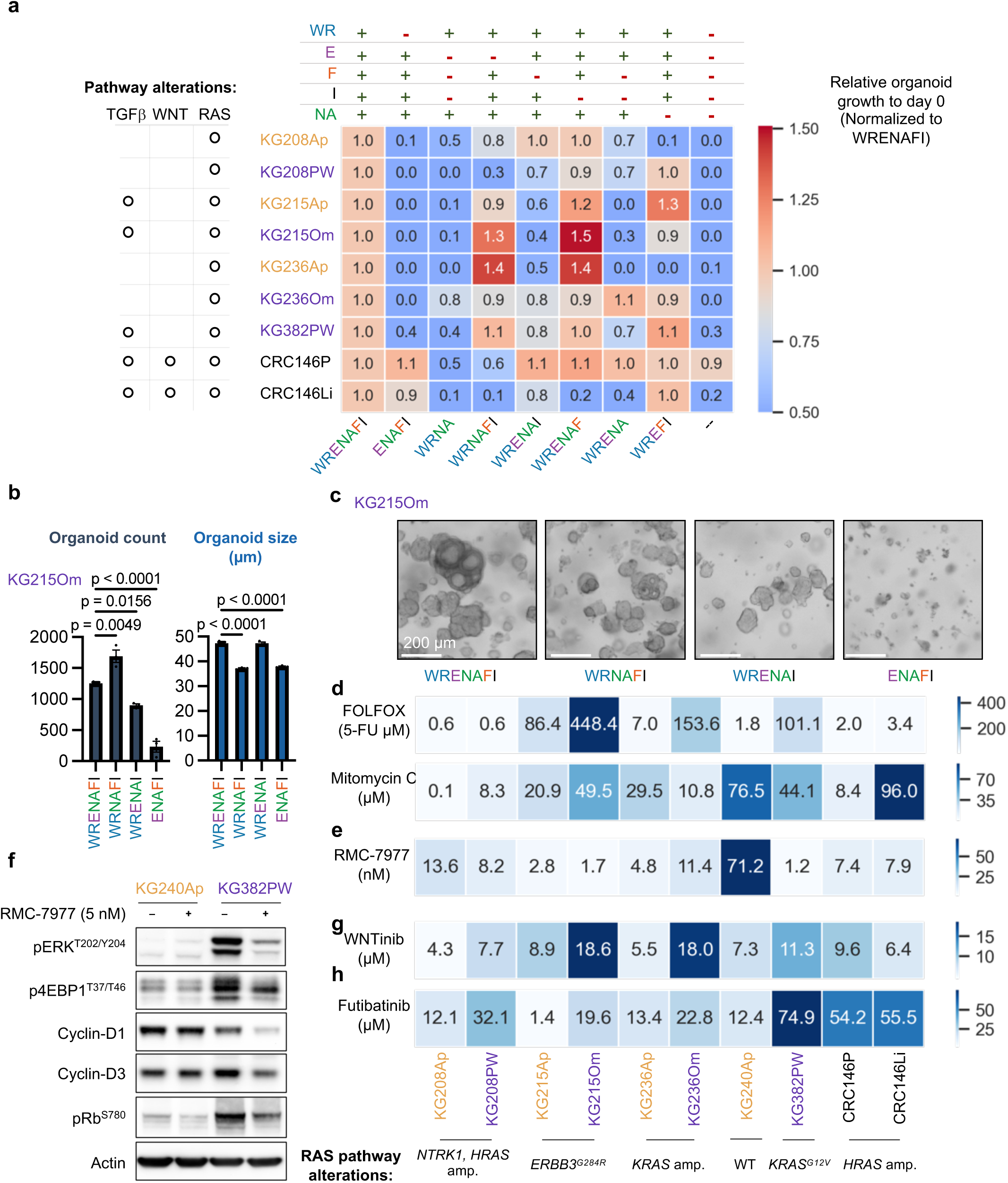
Analysis of niche-factor dependency reveals druggable vulnerabilities of AC. **a**, Heatmap indicating relative growth of the indicated AC and CRC organoid lines measured by CellTiter-Glo luminescence on day 7 relative to day 0 following seeding of 2000 single cells/well in the indicated media conditions and normalized to full media (WRENAFI) for each organoid line. n= 5-8 replicates per condition per line. W= Wnt, R = R-spondin, E =. Epithelial Growth Factor, N = Noggin, A= A83-01, F = Fibroblast Growth Factor 2, I= Insulin-like Growth Factor. ‘O’ indicates altered signaling. A pathway was considered altered when one or more genes in the signaling pathway were identified as having ‘sustained oncogenic” or “likely oncogenic” point mutations or copy number alterations (see Methods for specific genes), as assayed by MSK-IMPACT^82^ and identified by OncoKB^83^. **b**, Organoid counts (stemness) and size (proliferation) of KG215Om organoids after 7 days of growth in the indicated media. Mean ± s.e.m. n = 3, One-way ANOVA tests. **c**, Representative images of KG215Om PDOs in the indicated media conditions. **d, e, g, h**, Heatmaps showing the IC50 values of the indicated AC and CRC organoid lines seeded as 5000 single cells, grown for 5 days and then treated with **(d**) standard of care CRC cytotoxic chemotherapies (FOLFOX, mitomycin C), (**e**) RAS^MULTI^-(on) inhibitor RMC-7977, (**g**) multi-kinase/Wnt-signaling inhibitor WNTinib and (**h**) FGF2 inhibitor futibatinib. Values are of means of ≥3 replicates. **f**, Western blots showing activation status of RAS/MAPK pathway targets and regulators of cell cycle in KG240Ap (*KRAS^WT/WT^*) and KG382PW (*KRAS^G12V/WT^*) PDOs treated with RMC-7977 or DMSO control for 72hr.

### Niche-inspired targeted therapies for AC-PC

Our interrogation of the niche-dependencies of primary AC and AC-PC identified signaling pathways that are selectively upregulated in AC-PC, suggesting potential routes to targeting AC-PC using inhibitors of these pathways (**Figure 3a-c, S4c**, **S5**). We first asked whether AC organoids display sensitivity *ex vivo* to 5-fluorouracil + oxaliplatin (active components of the FOLFOX regimen) and mitomycin C, two standard of care chemotherapy regimens initially developed for CRC and commonly used to treat both C-PC and AC-PC^52–55^. We found tremendous heterogeneity in AC organoid sensitivity to FOLFOX and mitomycin C (**Figure 3d**). Notably, 2/3 AC-PC organoids had IC_50_ values that were 20-100 fold higher than that of their paired primary AC organoids. AC PDOs were similarly resistant to mitomycin C.

Next, since RAS pathway mutations are present in over 40% of AC and our finding that AC organoids are highly sensitive to withdrawal of MAPK ligands EGF, FGF and IGF (**Figure 3a, S5**), we tested the response of AC and CRC PDOs to the RAS-ON inhibitor RMC-7977, an analog of RMC-6236, currently in clinical trials^56,57^. We found that AC organoids that had alterations in the RAS pathway were sensitive RAS pathway inhibition (**Figure 3f, Table S1**). Notably, KG382PW, which harbors a *KRAS^G12V^*mutation, displayed the highest sensitivity, while KG240Ap, which retains wildtype RAS, displayed the least sensitivity to RMC-7977, consistent with prior observations in RAS WT/mut cancers^56,57^. To validate the on-target activity of RMC-7977in AC organoids, we performed western blot analysis of KRAS mutant (KG382PW) and KRAS wildtype (KG240Ap) PDOs and found that phosphorylation of ERK^T202/Y204^, 4EBP1^T37/46^ and Rb^S780^ phosphorylation as well as Cyclin D1 expression were downregulated after treatment with RMC-7977 in *KRAS^G12V^* mutant KG382PW, but not RAS-WT KG240Ap AC organoids (**Figure 3f**).

Beyond RAS inhibition, recent work has identified a novel RTK inhibitor, WNTinib, which effectively targets Wnt-dependent hepatocellular carcinoma^58^. Given the dependency of AC on both Wnt and RTK signaling for organoid formation and growth, we tested the ability of WNTinib to inhibit the growth of AC organoids. Both AC and CRC organoids were responsive to WNTinib, although metastasis organoids were less sensitive than their primary tumor counterparts (**Figure 3g**). Finally, FGF withdrawal from organoid media led to the most notable reduction in organoid formation and growth (**Figures 3a, c**), consistent with studies showing that FGF2 enhances intestinal stem cell function^59,60^. We therefore tested futibatinib, an FGFR inhibitor recently FDA-approved for the treatment of cholangiocarcinoma^61^. Similar to WNTinib, futibatinib was more effective in primary organoids compared to metastasis, but both were more sensitive to FGFR inhibition compared to CRC organoids (**Figure 3h**). Together, our preclinical testing using AC-PC organoids identify three inhibitors of MAPK/PI3K signaling — RMC-7977, WNTinib and futibatinib, as clinically actionable strategies for treating AC-PC, that are more effective *ex vivo* than current CRC-derived standard-of-care agents FOLFOX and mitomycin C.

### Preclinical efficacy of RMC-7977 and WNTinib in a diffuse peritoneal carcinomatosis mouse model

Given the *ex vivo* efficacy of RMC-7977 and WNTinib in inhibiting AC organoid growth, we sought to examine the ability of these agents to treat AC-PC in our clinically representative DPC model. In the clinic, where feasible patients typically undergo surgical debulking of PC and are then administered adjuvant chemotherapy to eliminate any residual cancer cells. Modeling this clinical scenario, we introduced *KRAS^G12V^*mutant KG382PW AC-PC organoids as diffuse peritoneal carcinomatosis, and then administered 5 daily doses of drug or vehicle control. While both RMC-7977 and WNTinib led to initial weight loss in the treated mice, all treated animals had regained weight by 6 weeks after treatment (**Figure S6a-c**), with post-mortem analysis of organs revealing no long-term adverse effects. The administration of RMC-7977 and WNTinib immediately after the transplantation of KG382PW in the peritoneum led to an immediate reduction of BLI signals and to a significant sustained decrease in tumor burden 8 weeks after the administration of the first dose (**Figure 4a-c**). *Ex vivo* analysis of KG382PW xenograft tumors harvested following three consecutive daily *in vivo* doses of RMC-7977 showed reduction in RAS signaling including reduction in MAPK signaling effectors pERK and p4EBP1 and cell-cycle activity, and increased apoptosis (**Figures 4d-e, S6d**). WNTinib treated tumors also displayed significant decrease in pERK staining (**Figure S6e**). Together, our results in our representative DPC *in vivo* model identify RMC-7977 and WNTinib as safe and effective, clinically actionable strategies for treating peritoneal carcinomatosis.

**Figure 4.**
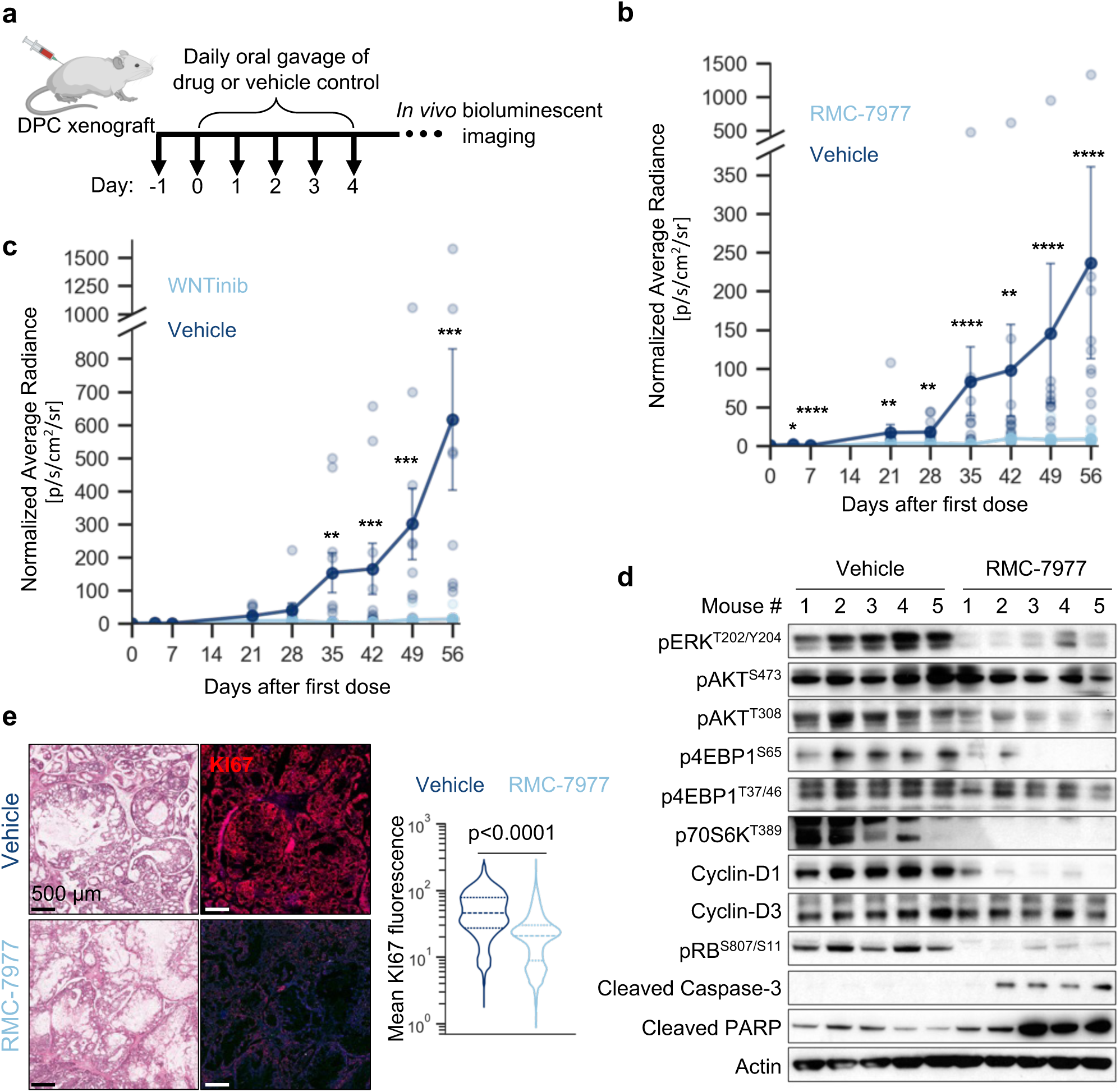
Preclinical efficacy of RMC-7977 and WNTinib in AC diffuse peritoneal carcinomatosis. **a**, Schematic of RMC-7977 and WNTinib experimental design. NSG mice were injected with 500,000 cells from KG382PW organoids to establish diffuse peritoneal carcinomatosis. Starting 24 hours after injection, animals were administered 5 daily doses of 25 mg/kg RMC-7977 or 25 mg WNTinib or vehicle controls. Peritoneal carcinomatosis growth was monitored using *ex vivo* bioluminescence imaging for 8 weeks. **b**, **c,** Normalized average radiance measured by *ex vivo* bioluminescence imaging following intraperitoneal injection of 500,000 organoids from KG382PW. Animals were dosed daily with (**b**) 25mg/kg RMC-7977 and (**c**) 25mg/kg WNTinib or corresponding vehicle via oral gavage for 5d. Mean ± s.e.m. (**b**) n = 10 vehicle, n = 9 RMC-7977. (**c**) n = 10 vehicle, n = 8 WNTinib. Significant differences are indicated by asterisk as calculated by two-sided Mann– Whitney *U* tests, *p < 0.05, **p < 0.01, *** p < 0.001, **** p < 0.0001. **c**, Representative H&E and Ki67 immunofluorescence staining (left) and quantification (right) of KG382PW diffuse peritoneal carcinomatosis xenograft tumors harvested from NSG mice treated with RMC-7977 or vehicle control as in (d). Violin plots show the distribution of KI67 mean fluorescence per DAPI positive (blue) nucleus. n = 163,110 total nuclei, 3 mice per group. Dashed line indicates the median and the dotted lines indicate the interquartile range. Two-sided Mann– Whitney *U* tests. **d**, Western blots showing activation status of RAS/MAPK pathway targets and regulators of cell cycle in KG240Ap (*KRAS^WT/WT^*) and KG382PW (*KRAS^G1V2/WT^*) PDOs treated with RMC-7977 or DMSO control for 72hr.

## Discussion

In this study, we established to our knowledge the first sustainable resource of patient-derived organoid models for studying AC and PC. We show that primary AC and AC-PC organoids not only retain intertumoral heterogeneity, cell morphology, and mutational profiles, but can also serve as platform for identifying actionable oncogenic mutations undetected in mucinous paucicellular patient tissue using routine next generation sequencing clinical diagnostic assays. Using our clinically representative patient organoid-derived DPC model, we show that metastasis-derived organoids retain the ability to colonize the peritoneum, unlike their patient-matched primary tumor counterparts, with scRNAseq of paired primary-metastasis organoids revealing reprogramming from mucinous intestinal states into dedifferentiated stem-like and endoderm-like states in peritoneal metastasis, concordant with the dedifferentiation observed in CRC liver metastasis^43^. Using scRNA-Seq, and functional growth factor withdrawal studies, we show that metastasis PDOs have cell-intrinsic upregulation of mitogenic signaling and are more resistant to withdrawal of extrinsic growth factors, which we hypothesize endows PC cells with the ability to survive in the relatively mitogen-depleted microenvironment of the peritoneum^62–64^.

Previous work has shown the organoids recapitulate responses to small molecules and immunotherapies, and that it can be predictive of patient response^29,65–68^. We show that AC PDOs have demonstrable resistance to CRC-inspired standard treatments, including FOLFOX and mitomycin C, underscoring the need to identify AC-specific therapeutic strategies. Building on our functional data and dependence on oncogenic signaling pathway activity, we identify several niche-inspired druggable dependencies of AC. We show that the inhibition of the FGFR, RAS and the Wnt pathways can effectively kill cancer cells *ex vivo*. We further show that the RAS-ON inhibitor RMC-7977 and WNTinib can reduce DPC growth *in vivo*. RMC-7977 was highly efficacious as a single agent against AC-PC *in vivo*, with efficient on-target inhibition of pERK and p4EBP1 which have been shown to correlate with MAPK-inhibition associated apoptosis^69,70^. In contrast, in prior studies in breast, colorectal and non-small cell lung cancer, dual inhibition MAPK and PI3K signaling by targeting AKT/ERK or KRAS^G12C^/mTORC1 was required to induce 4EBP1 phosphorylation, effective inhibition of cap-dependent translation, and apoptosis^69,70^. These data indicate that AC might be particularly dependent on high RAS activity, as suggested by the high frequency of oncogenic MAPK mutations in AC vs CRC (*KRAS* (58% vs 44%) and *GNAS* (33% vs 5%))^23^. The increased sensitivity of RAS^mut^ organoid lines to RMC-7977 (**Figure 3e**), underscores the importance of correctly identifying patients with KRAS mutant tumors likely to benefit most from KRAS targeting agents, which is enabled by enrichment of cancer cells during organoid culture (**Figure 1e, f**). Our work paves the way for the future development of clinical trials for PC using *KRAS* or other MAPK inhibitors in the adjuvant setting, to target residual disease remaining after cytoreductive surgery, decreasing peritoneal metastasis relapse and prolonging overall survival. More broadly, we show that our appendiceal cancer organoid and organoid-derived DPC models can serve as a powerful preclinical platform for identifying actionable mutations, and testing novel therapeutic strategies to nominate candidates for clinical testing in patients whose tumors are resistant to standard treatment.

Our DPC model recapitulates the distinct fat-tropism of PC-metastatic tumors. While previous studies have shown that fatty tissue offers a distinct tumor microenvironment than other common metastatic sites, such as the liver or bone marrow^71–73^, they have been limited to the use of *in vitro* cultured adipocytes to interrogate cancer cell-adipocyte interactions. Our organoid:DPC model could be leveraged to understand the unique microenvironment of peritoneal tumors *in vivo*. In addition to AC, peritoneal metastasis is common in ovarian, gastric and colorectal cancers and is frequently associated with aggressive disease and poor survival^74–80^. Our *in vivo* model recapitulates diffuse abdominal tumors characteristic of PC and can serve as a valuable disease-relevant framework for preclinical and mechanistic studies of PC across cancer types.

In sum, our work addresses a significant unmet clinical need by establishing an AC and AC-PC organoid biobank and mouse models that faithfully recapitulate clinical, genetic and morphological features of diffuse peritoneal carcinomatosis. We highlight the use of organoids as diagnostic platforms for identifying overlooked actionable oncogenic mutations in paucicellular mucinous AC tumors. We demonstrate the limitations of CRC-derived treatments in targeting AC, and identify RTK-targeting agents as effective therapeutic agents to treat PC *in vivo*.

## Methods

### Tissue processing for organoid derivation

Patients undergoing surgical resection of appendiceal cancer primary and/or metastatic disease at MSKCC were identified by chart review, and those who had signed pre-procedure informed consent to MSK IRB protocols #06-107, #12-245, #14-244 and #22-404 for biospecimen collection were selected for this study. Freshly resected surgical tissue in surplus of clinical diagnostic requirements was processed to generate organoids. Portions were also fixed in formalin and embedded in paraffin. Tissue was generally processed within 1 h of surgical resection. Archival formalin-fixed, paraffin-embedded (FFPE) clinical tissue blocks for immunostaining were identified by database search and chart review. Tissue processing and histopathological data interpretation were overseen by an expert gastrointestinal pathologist (J.S.). Primary and metastatic appendiceal tumors were processed for organoid derivation as previously described^43^. Briefly ∼200 mg freshly resected tumor tissue was collected in tissue collection media (Advanced DMEM/F12; (Thermo Fisher Scientific), GlutaMAX (2 mM, Thermo Fisher Scientific), HEPES (10 mM, Thermo Fisher Scientific), *N*-acetyl-L-cysteine (1 mM, Sigma-Aldrich), B27 supplement with vitamin A (Thermo Fisher Scientific) supplemented with primocin (100 μg ml^−1^, InvivoGen), plasmocin (50 μg ml^−1^, InvivoGen), penicillin–streptomycin (100 μg ml^−1^, Thermo Fisher Scientific), Amphotericin B (2.5 μg ml^−1^, Cytiva), nystatin (250 U ml^−1^, Millipore Sigma). Specimens were placed into a 15 cm Petri dish using sterile forceps and washed three times with DPBS (Thermo Fisher Scientific) supplemented with the above antibiotics and chopped into 2-mm fragments. Tumor fragments were transferred into a gentleMACS type C tube (Miltenyi) pre-filled with 5 ml of WRENAFI medium supplemented with DNase I (100 U ml^−1^, Millipore Sigma) and a proprietary cocktail of tissue digestion enzymes (Tumour Dissociation Kit, Miltenyi). Tumors were digested using the gentleMACS Octo Dissociator according to the manufacturer’s 37C_h_TDK_1 protocol for 30-60 minutes. The digested tissue suspension was filtered through a 100-μm strained and pelleted for 5 min at 500g and 4 °C. For MCC samples, if a mucinous layer remained visible after digestion, the pellet was treated with 10 mL TrypLE (Thermo Fisher Scientific) for 10 minutes at 37 °C. If red blood cells were visible in the cell pellet, 5 mL of ACK lysis buffer (Lonza) would be added to the pellet and incubated at room temperature for 5 minutes. CRC organoids were established previously as described^43,81^.

### Organoid culture and assays

AC and CRC organoids were cultured in WRENAFI (Advanced DMEM/F12 (AdDF12; Thermo Fisher Scientific), GlutaMAX (2 mM, Thermo Fisher Scientific), HEPES (10 mM, Thermo Fisher Scientific), N-acetyl-l-cysteine (1 mM, Sigma-Aldrich), B27 supplement with vitamin A (Thermo Fisher Scientific), primocin (100 μg ml^-1^, InvivoGen), EGF (50 ng ml^-1^, Peprotech), Noggin (50X, IPA), A83-01 (500 nM, Sigma-Aldrich), FGF2 (50 ng ml^-1^, Peprotech), IGF-I (100 ng ml^-1^, Peprotech), 1% R-pondin1-conditioned medium (media collected from HEK293 cell lines expressing recombinant R-spondin 1), NGS-Wnt (0.5 M, ImmunePrecise N001). Cells processed as described above were suspended at 2,000 cells per 40 μL and plated in 40 μL domes of 1:1 Matrigel:WRENAFI mixture (Corning) in suspension culture plates. After Matrigel domes solidified at 37 °C for 30-60 minutes, WRENAFI supplemented with Y-27632 (MedChemExpress) was added to the well.

For passaging, the Matrigel was depolymerized by adding 3mM EDTA in DPBS, mechanically disrupting the Matrigel domes with a p1000 pipette, and incubating the culture plate at 4 °C for 30-60 minutes. The cells suspension was centrifuged for 5 min at 150*g*/4 °C and washed with DPBS supplemented with, GlutaMAX (2 mM), HEPES (10 mM) and penicillin–streptomycin (100 IU ml−1, 0.1 mg ml−1, ThermoFisher Scientific). 3-5 day old organoids were harvested from Matrigel and cryopreserved in BAMBANKER™ serum-free cell freezing medium (FUJIFILM Irvine Scientific). For niche factor withdrawal assays, organoids were collected from Matrigel 3mM EDTA in DPBS then dissociated into single cells using TrypLE (Thermo Fisher Scientific) for 5–10 min at 37 °C, serially filtered through a 40-μm and 20-μm cell strainers to generate single cells suspensions. Cells were plated at 5000 cells/ 40 μL Matrigel and cultured in specified media supplemented with Y-27632. Media was changed every 3 days. Cell viability was measured using CellTiter-Glo assays (Promega) seven days after seeding. For the organoid formation assays, cells were seeded as above in 12-well suspension plates and imaged seven days after seeding. Imaging and luminescence reading were done using a BioTek Cytation 5 Cell Imaging Multimode Reader. Organoid numbers were quantified using the Agilent BioTek Gen5 data collection and analysis software. For drug sensitivity assays, organoids were processed into single cells as above and seeded in 96-well plates at 2000 cells/40 μL Matrigel/well with 50 μL WRENAFI. Organoids were grown for 4 days before addition of indicated drugs or vehicle. 5-FU (Selleckchem), Oxaliplatin (MedChemExpress) – the FOLFOX cocktail was prepared at a ratio of 25:1 5-FU:Oxaliplatin; mitomycin C (Fisher Scientific); Futibatininb (Selleckchem); RMC-7977 was a kind gift from Neal Rosen (Memorial Sloan Kettering Cancer Center); WNTinib was a kind gift from Arvin Dar (Memorial Sloan Kettering Cancer Center). Three days after adding the drugs, cell viability was measured using CellTiter-Glo using a BioTek Cytation 5 Cell Imaging Multimode Reader. IC50 values were calculated by fitting normalized percent viability to a nonlinear regression curve.

### Immunostaining

Organoids were harvested from Matrigel as above and resuspended in a 250 μL solution of 10% 10X PBS, 2% 1 M NaOH and 88% Collagen I (Corning). After solidifying for 30 min at 37 °C, the collagen discs were fixed in 4% paraformaldehyde, followed by paraffin embedding using standard protocols. Human tissues were fixed in formalin and mouse tissues were fixed in 4% paraformaldehyde for 24 hours before paraffin embedding, sectioning (5 μm). Immunohistochemistry staining was performed by the Molecular Cytology Core Facility of Memorial Sloan Kettering Cancer Center using a Discovery XT processor (Ventana Medical Systems) using standard automated protocols, SYP (clone Snp-88, BioGenex) and CDX2 (clone CDX2-88, BioGenex) antibodies. For immunofluorescence staining, slides were dewaxed with Histo-Clear® (VWR) at room temperature for 10 min rehydrated using standard protocols, processed through heat-induced antigen retrieval using universal HIER antigen retrieval reagent (Abcam), washed three times with PBS, blocked using 10% Normal Goat Serum (ThermoFisher) at room temperature for 20 min, washed 3 times with PBS and stained with primary antibodies overnight at 4 °C. Slides were then washed 3 times with PBS and stained with secondary antibodies for 60 min. One drop of VECTASHIELD® Antifade Mounting Medium with DAPI (Vector Laboratories) mounting media was then added to the slides and No.1 ½ cover glasses (Millipore Sigma) were attached. Primary antibodies used for staining were E-cadherin (clone 24E10, Cell Signaling, 1:1600), Chromogranin A (clone 24E10, Novus Biologics, 1:500), Cytokeratin 20 (clone EPR1622Y, Abcam, 1:100), Mucin 2 (clone CCP58, Novus Biologics, 1:1000), CDX2 (clone EPR2764Y, Abcam, 1:200), Cytokeratin 7 (clone OV-TL12/30, Novus Biologics, 1:750), KI67 (clone SP6, Abcam, 1:100) and KI67 (clone SP6, Abcam, 1:100). Secondary antibodies used for staining were Alexa Fluor 488 goat polyclonal anti-mouse and Alexa Fluor 594 goat polyclonal anti-rabbit IgG (H+L) (Thermo Fisher Scientific). Slides were scanned on a Pannoramic Scanner (3DHistech, Budapest, Hungary) using a 20x/0.8NA objective. For IF quantification, regions of interest were drawn manually using QuPath to include only tumor cells and mean Ki67 or pERK signal per nucleus was used.

### Targeted next generation sequencing by MSK-IMPACT

DNA was isolated from organoids using the PicoPure™ DNA Extraction kit (Thermo Fisher Scientific) according to the manufacturer’s protocol. Tumor and organoid DNA was sequenced using targeting DNA sequencing using the MSK Integrated Mutation Profiling of Actionable Cancer Targets (MSK-IMPACT) platform as previously described^82^. Organoid sequencing data underwent variant calling through the ARGOS pipeline (https://github.com/mskcc/argos-cwl/). Oncogenic or likely oncogenic mutations were identified using the OncoKB^83^. Mutations that were discordant between tumor-organoid pairs in frequent driver genes of appendiceal cancer were selected for further investigation. This process involved reviewing the sequencing BAM file of the discordant tumor sample to determine if any reads supporting the mutation were present. Genes sequenced through MSK-IMPACT in the WNT, TGFβ and RAS pathways are: WNT: *SFRP1, SFRP2, SFRP3, SFRP4, SFRP5, WIF1, DKK1, DKK2, DKK3, DKK4, RNF43, LPR5, FZD, LRP6, GSK4B, CTNNB1, APC, AMER1, AXIN1, AXIN2*; TGFβ: *TGFBR1, TGFBR2, SMAD2, SMAD3, SMAD4, ACVR2A, ACVR2B*; RAS: *ROS1, KIT, RET, MET, FGFR2, ALK, FLT3, NTRK2, FGFR3, FGFR1, ERBB2, NTRK1, ERBB4, EGFR, FGFR4, IGF1R, ERBB3, PDGFRA, CBL, ERRFI1, PTPN11, SOS1, NF1, NRAS, RIT1, HRAS, KRAS, RASA1, ARAF, RAF1, BRAF, MARK2K1, MAP2K2, MAPK1, RAC1*^82,84^.

### Diffuse peritoneal carcinomatosis xenograft model

Female NSG (*NOD.Cg-Prkdc^scid^Il2rg^tm1Wjl^/SzJ*, 005557) mice were obtained from the Jackson Laboratory and were transplanted at 5 weeks of age. Mice were maintained in a specific-pathogen-free (SPF) facility under a 12 h–12 h light–dark cycle under controlled temperature and humidity and maintained on an amoxicillin diet. Organoids were transduced with pLenti-PGK-tdTomato-AkaLuc^43^, and selected by culture in WRENAFI media containing 60 μg/mL Geneticin™ (ThermoFisher Scientific) following by FACS sorting for tdTomato positive cells using a 130 μm (SH800S SONY Sorter). For the initial optimization experiments, 500,000 cells from MSK107Li CRC organoids in WRENAFI were injected intraperitoneally in either 0:1, 1:1 or 3:1 100 μL WRENAFI:Matrigel solution. To aid the diffuse spread of the injected cell suspension, the mouse abdomen was continually massaged for 30-60 seconds following injection. For all subsequent experiments, 500,000 cells of the indicated organoids stably transduced with pLenti-PGK-tdTomato-AkaLuc were injected in 100 μL 3:1 WRENAFI:Matrigel solution. For *ex vivo* bioluminescent imaging, mice were injected IP with 100 μL of 3.3 mM TokeOni (Sigma-Aldrich) in sterile saline and imaged 15 min after substrate administration using an IVIS Spectrum Xenogen instrument (Caliper Life Sciences). BLI images were analyzed using the Living Image software v.2.50.

### Single-cell RNA-Sequencing (scRNA-Seq)

#### Sample preparation for scRNA-seq

Organoids were harvested from Matrigel 3mM EDTA in DPBS then dissociated into single cells using TrypLE (Thermo Fisher Scientific) for 5–10 min at 37 °C and serially filtered through a 40-μm then 20-μm cell strainers to generate single cell suspensions. For multiplexing, cells were incubated with TotalSeq-B0251 or TotalSeq-B0252 hash-tagging antibodies in FACS buffer (10 mM HEPES, 0.1 mM EDTA, 0.1% FBS) for 45 min then washed 3X with FACS buffer. Cells were then incubated with DAPI in FACS buffer sorted to enrich for live cells then equal numbers of cells from each of two samples were combined into a single suspension. scRNA-seq was performed on the Chromium instrument (10x Genomics) according to the 3′ RNA v3.1 user manual as previously described^43^.

#### Integration with scVI/scANVI

A number of integration methods were tested on the data set and evaluated using the scIB pipeline fast metrics^85^ which measure both batch correction and preservation of biological variance. The overall best performer was scANVI^86^ a semi-supervised variant of scVI^87^, a deep-learning method that uses a variational autoencoder network. First, a hyperparameter grid search was performed and a scVI model with 3 hidden layers of 64 nodes per layer and 40 latent dimensions selected, with dispersion allowed to differ between batches. The default number of epochs with early stopping based on reconstruction loss validation was used for all training. The trained scVI model was then used to generate a scANVI model supplied with sample type labels (primary tumor vs. metastasis). The scANVI model was trained for an additional 71 epochs after which it showed no further improvement.

#### GSEA and gene signature scores

The CellBender-denoised count matrix was normalized to median library size and log-transformed. Differentially expressed genes were calculated between primary-metastasis matched pairs using the R package MAST (v.1.16.0) and ranked each DEG by −log[P] × log[fold change] value. GSEA was performed using the Python package gseapy (v.1.1.3) using relevant gene sets from the literature, our newly derived hotspot modules as well as Hallmark and KEGG. To generate all gene signature scores, we used Scanpy (v.1.9.8) score_genes function with z-normalized expression data as input.

### *In vivo* drug studies

500,000 KG382PW PDOs were orthotopically introduced into NSG mice as diffuse peritoneal carcinomatosis xenografts as above. Bioluminescent imaging was performed immediately after intraperitoneal injection and abdominal massage, and animals were randomized and assigned to either vehicle control or drug treatment groups. 100 μL of RMC-7977, WNTinib, or corresponding vehicle were administered at 25 mg/kg through oral gavage for 5 consecutive days starting 24 hours following injection. For *in vivo* delivery, RMC-7977 was reconstituted in 10% DMSO, 20% PEG-400 (Sigma-Aldrich), 10% Solutol® (Sigma-Aldrich) and 60% sterile water, and WNTinib was reconstituted in 25% 1:1 Solutol®:Ethanol solution and 75% sterile water. Experimental end point was reached when tumor size exceeded >10% of body mass, or when animals showed any signs of respiratory distress or illness, such as hunched posture, failure to groom or greater than 10–15% weight loss. Mice were euthanized at the humane timepoint, and normal and tumor tissue was collected for fixation in 4% paraformaldehyde and subsequent immunostaining, or flash frozen in liquid nitrogen for proteomic analysis.

## Acknowledgements

This work was supported by grants from the Dalton Appendix Foundation (K.G.), National Institutes of Health (U2CCA233284 (K.G.), U54CA209975 (K.G.), R37CA266185 (K.G.), K08CA230213 (K.G.), P30CA008748 (MSKCC)), Howard Hughes Medical Institute Gilliam Fellowship (A.M.), National Science Foundation GRFP (A.M.), Damon Runyon Clinical Investigator Award (K.G.), Burroughs Wellcome Career Award for Medical Scientists (K.G.), AACR NextGen Grant for Transformative Cancer Research (K.G.), Stand Up to Cancer Convergence 3.1416 Award (K.G.), Pershing Square Sohn Prize (K.G.), Starr Cancer Consortium (K.G.), Josie Robertson Investigator Award (K.G.), Gerry Metastasis and Tumor Ecosystems Center grants (Q.J. and K.G.).

## Declaration of interests

K.G. is listed as an inventor on US patent 11,464,874, and US provisional patent applications 63/478,809 and 63/478,829 on targeting L1CAM to treat cancer, submitted by MSKCC. J.S. is a consultant for Paige AI.

## Supplementary Tables

**Supplementary Table 1. Patient metadata, clinical information and mutations of AC patient-derived organoid biobank.**

---

**Figure S1.**
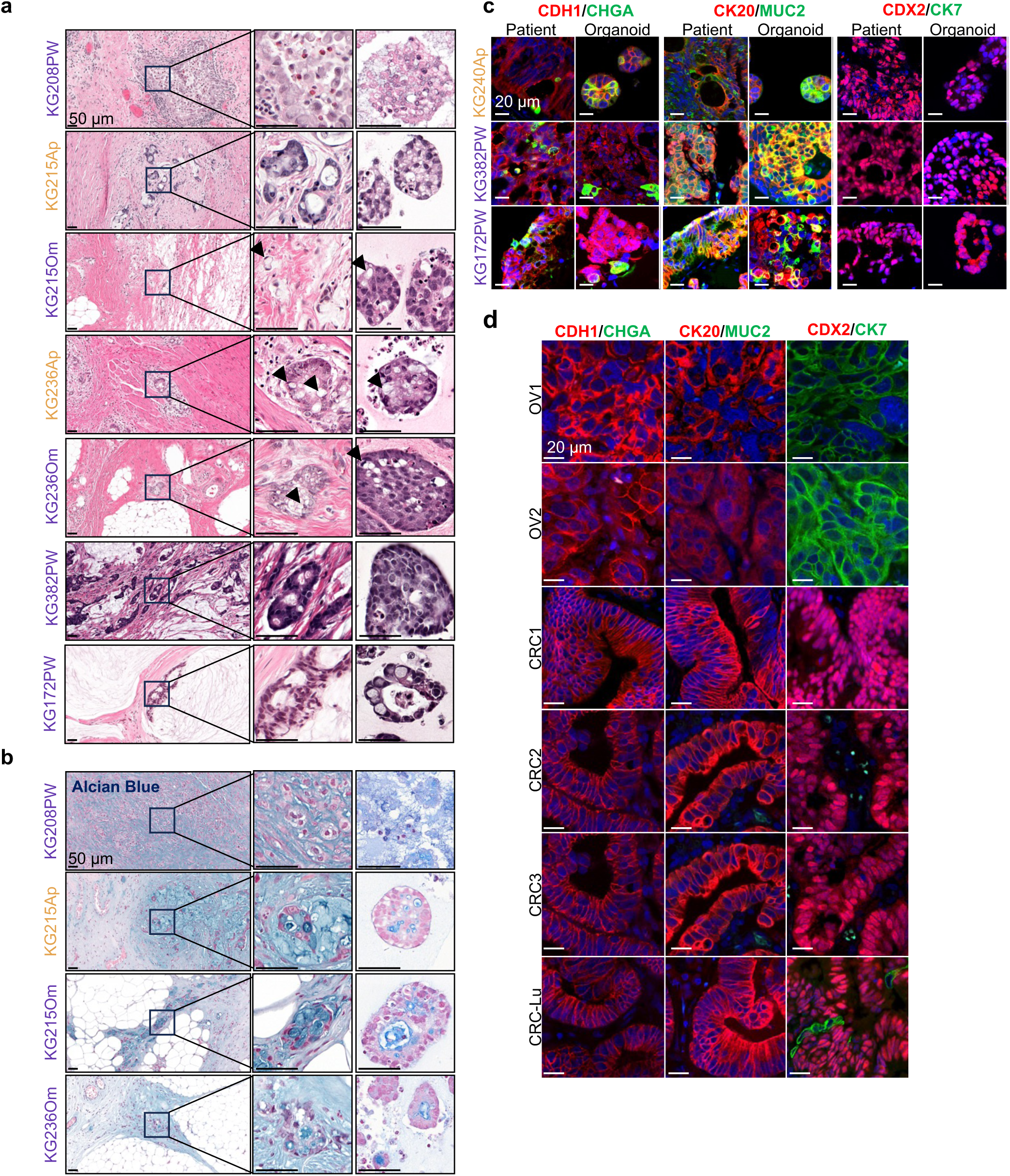
Histopathology and immunophenotype of AC tumors and organoids. a,b, Representative images of **(a)** H&E and **(b)** alcian blue staining of PDOs and corresponding parental tumors. **c,d,** Immunofluorescence images of CDH1 (red) & CHGA (green); KRT20 (red) & MUC2 (green); CDX2 (red) & CK7 (green) representing the immunophenotype of three primary (yellow label)-metastasis (purple label) **(c)** matched MC AC PDOs and corresponding parental tumors and **(d)** patient ovarian tumor (OV1-3), primary CRC tissue (CRC1-3) and lung-metastasis from CRC primary (CRC-Lu). Left: CDH1 (red) & CHGA (green); Center: KRT20 (red) & MUC2 (green); Right: CDX2 (red) & CK7 (green). Nuclei are counterstained with DAPI (blue). DAPI (blue) was used as a nuclear stain. Representative images of H&E staining of PDOs and corresponding parental tumors.

**Figure S2.**
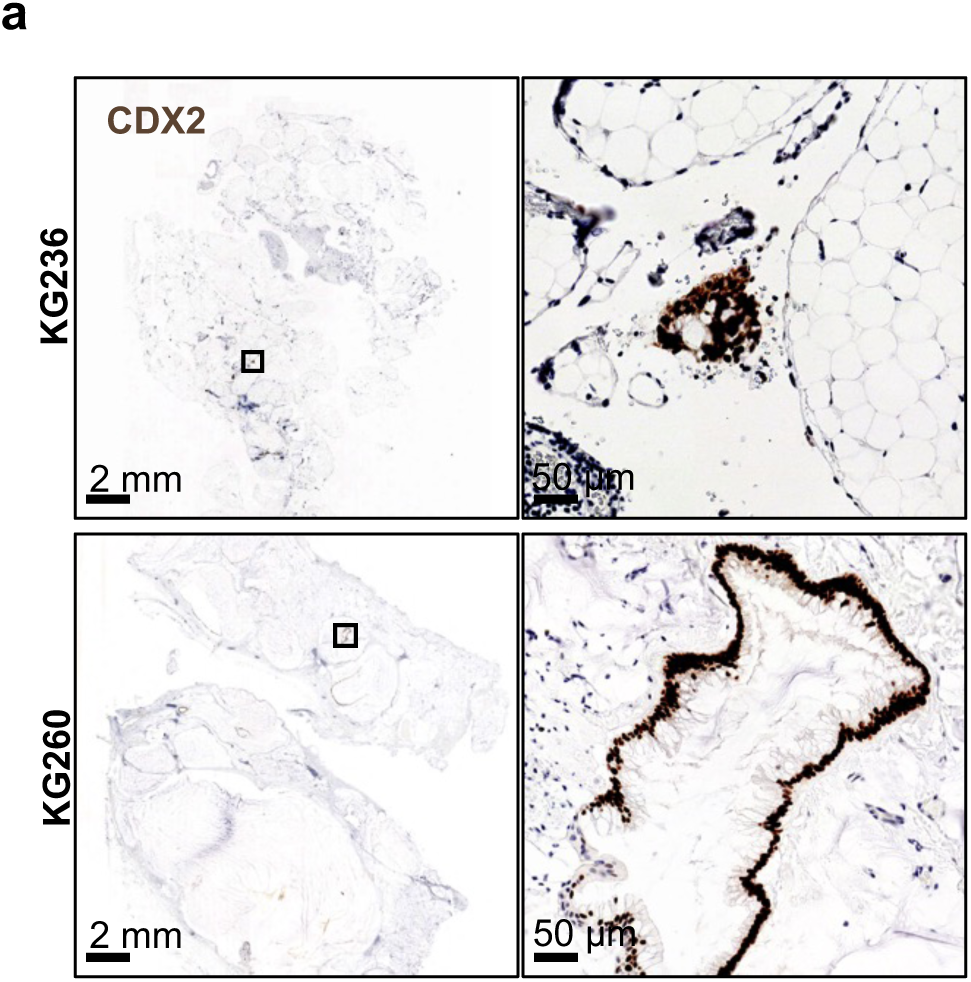
Oncogenic mutations are underrepresented in paucicellular AC tumors. a, CDX2 IHC staining of KG236 and KG260 clinical samples showing the low cellularity of AC tumors.

**Figure S3.**
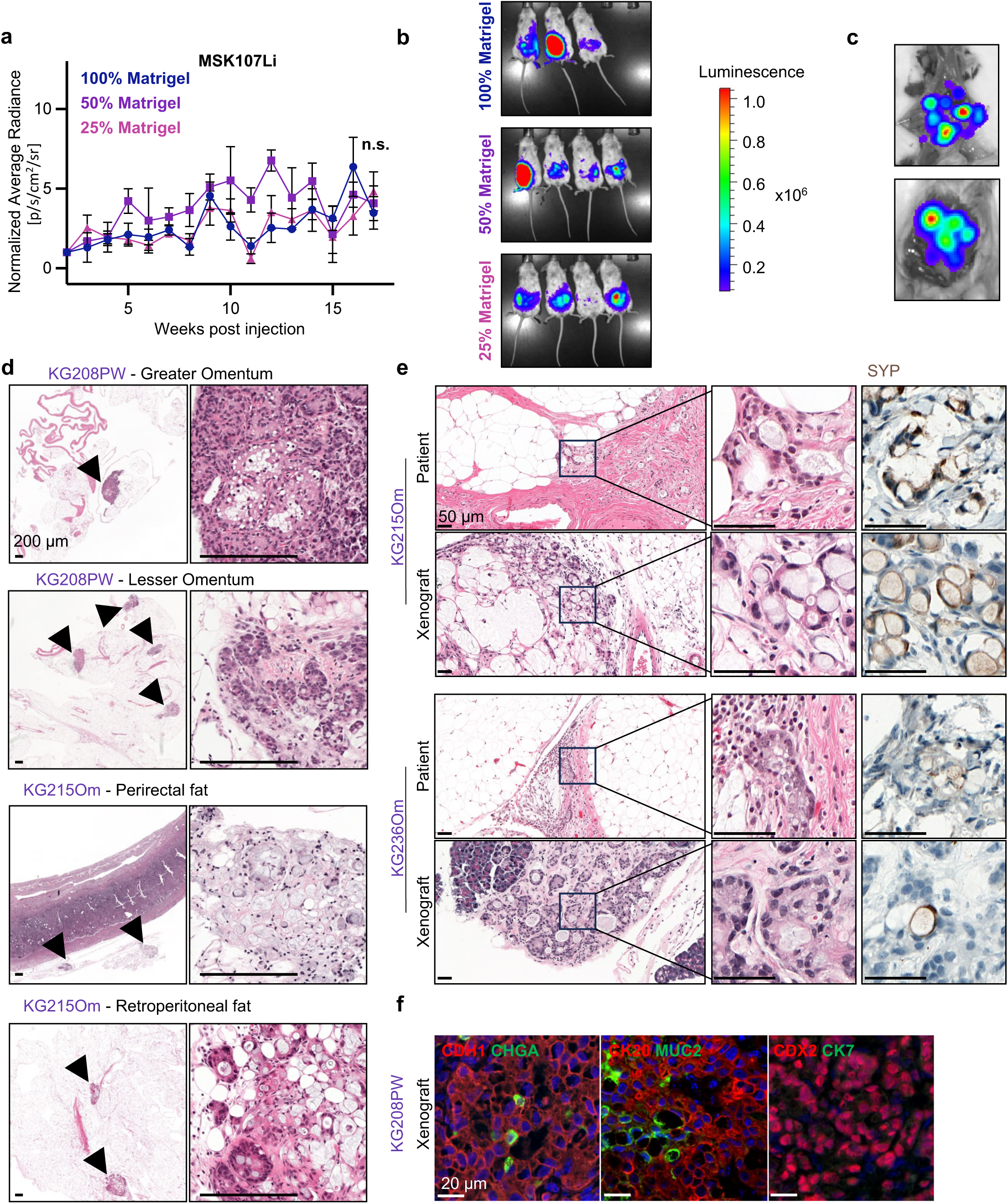
Optimization and characterization of peritoneal carcinomatosis xenograft model. a, Normalized average radiance measured by biweekly *ex vivo* bioluminescence imaging following intraperitoneal injection of 500,000 cell from MSK107Li organoids in either 25, 50, or 100% Matrigel. Mean ± s.e.m. n = 3 100% Matrigel, n = 4 50% Matrigel, n = 3 25% Matrigel. **b,** *Ex vivo* bioluminescence images of mice from **(a)** showing tumor growth 14, 15, 16 and 17 weeks post-intraperitoneal injections. **c**, Representative bioluminescence image showing diffuse peritoneal carcinomatosis post-mortem 10 weeks following intraperitoneal injection of 500,000 cells from KG215Om organoids. **d,** Representative H&E images of PDO-xenograft tumors in the specified fatty tissue 10 weeks after intraperitoneal injection of 500,000 organoids from the specified PDO line. **e, f,** Representative images of **(d)** H&E and IHC staining of the neuroendocrine marker synaptophysin (SYP) and **(e)** immunofluorescent staining of PDO-xenograft tumors 10 weeks after intraperitoneal injection of 500,000 organoid cells from the specified PDO line and the corresponding parental tumor. Left: CDH1 (red) & CHGA (green); Center: KRT20 (red) & MUC2 (green); Right: CDX2 (red) & CK7 (green). Nuclei are counterstained with DAPI (blue). DAPI (blue) was used as a nuclear stain.

**Figure S4.**
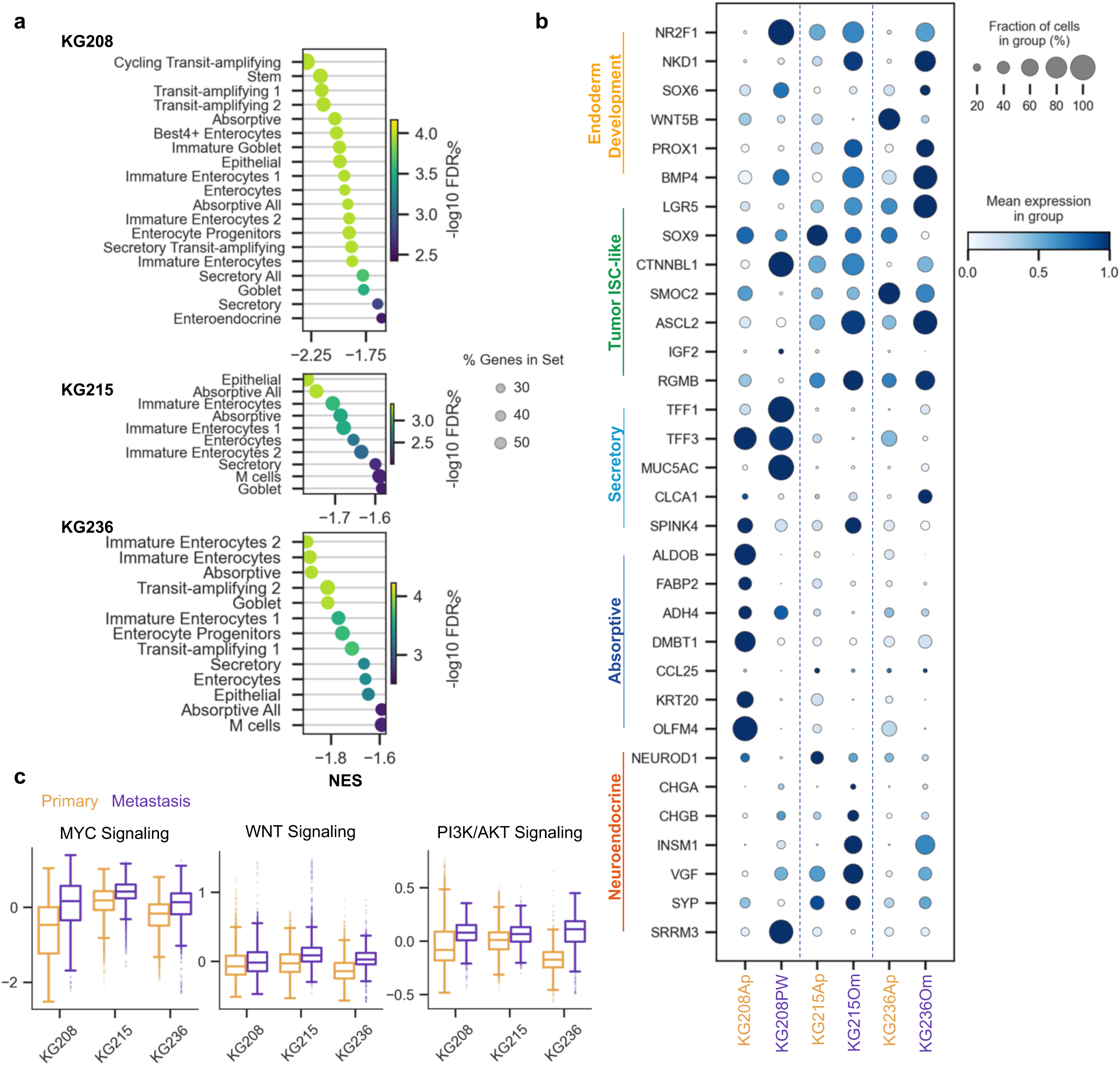
Single-cell transcriptomics identifies intrinsic metastatic capacity. a, GSEA normalized enrichment score (NES) of gene sets enriched in metastasis organoids as compared to the matched primary. **b,** Normalized gene expression of indicated genes within each sample. **c,** Average score for Hallmark gene sets calculated per cell.

**Figure S5.**
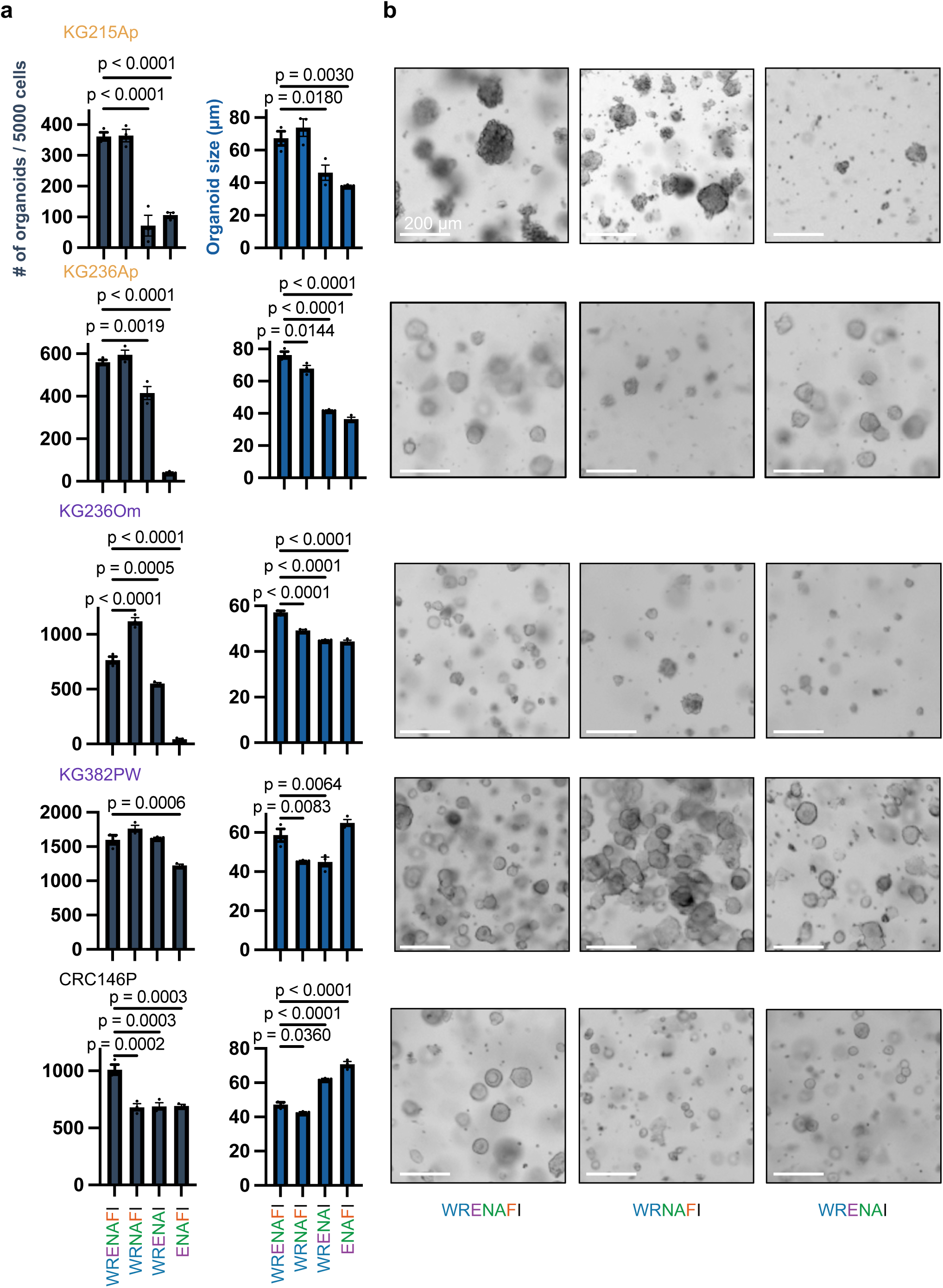
Effect of niche factor deprivation on stemness and proliferation. a, Organoid counts and size of indicated organoids after 7 days of growth in specified media. Organoids were dissociated and 5000 single cells were seeded and cultured +/-Wnt/R-spondin, FGF2 or EGF. Mean ± s.e.m. of n = 3 replicates, statistical analysis was performed using one-way ANOVA. **b,** Brightfield images of PDOs from **(a)**.

**Figure S6.**
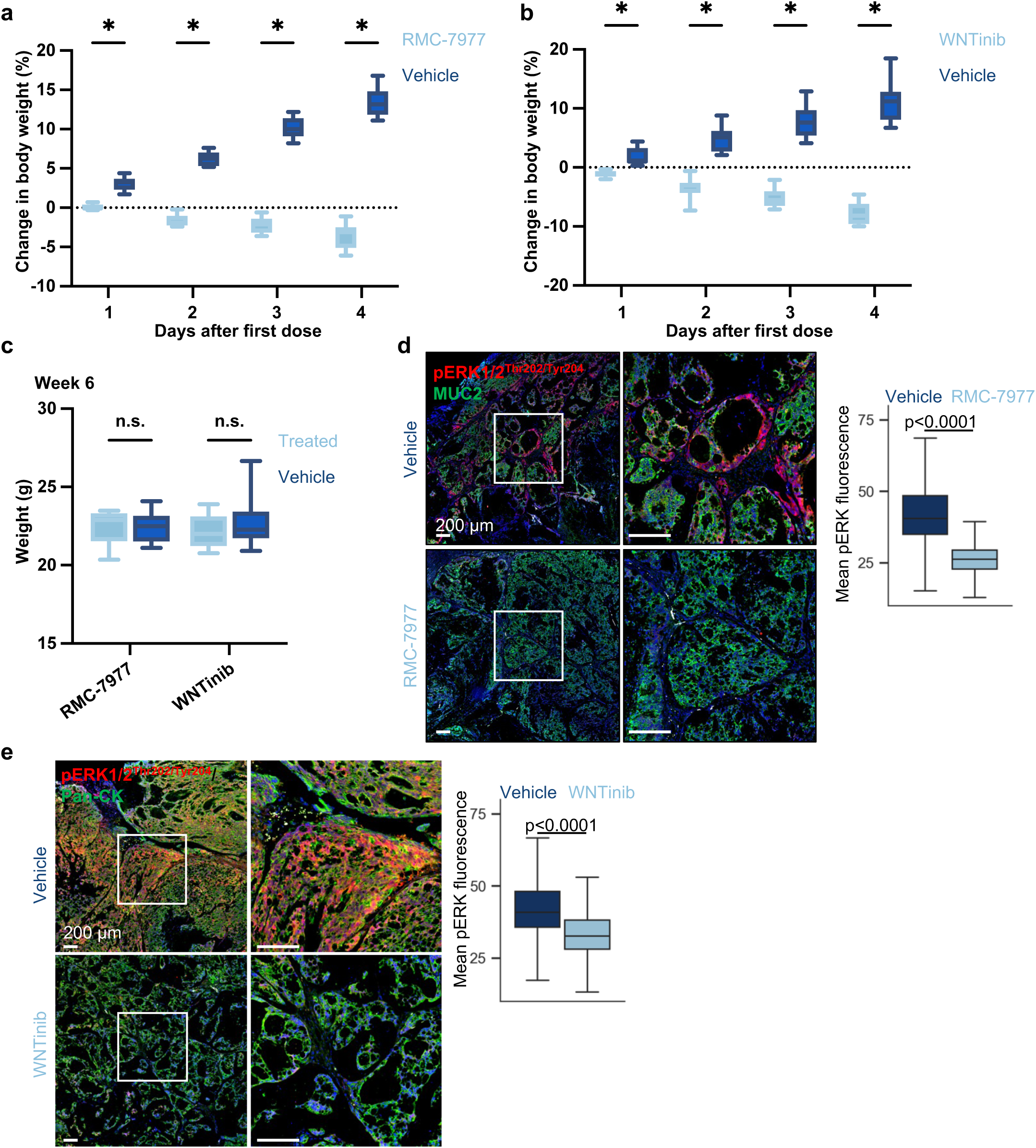
Efficacy and toxicity of WNTinib and RMC-7977 *in vivo*. a,b, Percentage change in body weight compared to D0 of mice from **(a)** Figure 4b and **(b)** Figure 4c. **c,** Body weights of NSG mice from Figures 4b**, c** six weeks after administration of the first dose of WNTinib, RMC-7977 or vehicle control. **d,** Immunofluorescence images of pERK1/2^Thr202/Tyr204^ in KG382PW peritoneal tumors from mice treated with RMC-7977 via oral gavage at 25mg/kg or vehicle control for 3d. Box plots show the distribution of pERK1/2mean fluorescence per nucleus. DAPI (blue) was used as a nuclear stain. n = 283,559 nuclei from 3 mice per group. Significant difference calculated by two-sided Mann–Whitney *U* tests. **e,** Immunofluorescence images of pERK1/2^Thr202/Tyr204^ in KG382PW peritoneal tumors from mice treated with daily WNTinib via oral gavage at 25mg/kg or vehicle control for 3d. Box plots show the distribution of pERK1/2mean fluorescence per nucleus. DAPI (blue) was used as a nuclear stain. n = 384,982 nuclei from 3 mice per group. Significant difference calculated by two-sided Mann–Whitney *U* tests.

## Key resources table

| REAGENT or RESOURCE | SOURCE | IDENTIFIER |
| --- | --- | --- |
| **Antibodies** |  |  |
| Rabbit polyclonal anti-phospho-MAPK (Thr202/Tyr204) | Cell Signaling Technology | Cat#9101<br>RRID:AB_331646 |
| Rabbit monoclonal anti-E-cadherin (clone 24E10) | Cell Signaling Technology | Cat#3195<br>RRID: AB_2291471 |
| Mouse monoclonal anti-MUC2 (clone CCP58) | Abcam | Cat# NBP2-25221<br>RRID:AB_2736858 |
| Rabbit monoclonal anti-CDX2 (clone EPR2764Y) | Abcam | Cat# ab76541<br>RRID:AB_1523334 |
| Rabbit monoclonal anti-Cytokeratin 20 (clone EPR1622Y) | Abcam | Cat# ab76126<br>RRID:AB_1310117 |
| Rabbit monoclonal anti-KI67 (clone SP6) | Abcam | Cat# ab16667<br>RRID:AB_302459 |
| Mouse monoclonal anti-CK7 (clone OV-TL12/30) | Novus Biologicals | Cat# NBP2-44814<br>RRID:AB_3309048 |
| Mouse monoclonal anti-keratin (clone C11) | Cell Signaling Technology | Cat# NBP2-44814<br>RRID:AB_490860 |
| Alexa Fluor 488 goat polyclonal anti-mouse IgG (H+L) | Thermo Fisher Scientific | Cat# A-11001<br>RRID:AB_2534069 |
| Alexa Fluor 594 goat polyclonal anti-rabbit IgG (H+L) | Thermo Fisher Scientific | Cat# A11012<br>RRID:AB_2534079 |
| Rabbit monoclonal anti-phospho-MAPK (Thr202/Tyr204) (clone D13.14.4E) | Cell Signaling Technology | Cat#4370<br>RRID:AB_2315112 |
| Mouse monoclonal anti-phospho-AKT (Ser473) (clone D9E) | Cell Signaling Technology | Cat#4060<br>RRID:AB_2315049 |
| Rabbit monoclonal anti-phospho-Akt (Thr308) (clone C31E5E) | Cell Signaling Technology | Cat#2965<br>RRID:AB_2255933 |
| Rabbit polyclonal anti-phospho- 4E-BP1 (Ser65) | Cell Signaling Technology | Cat#9451<br>RRID:AB_330947 |
| Rabbit polyclonal anti-phospho-4E-BP1 (Thr37/46) | Cell Signaling Technology | Cat#9459<br>RRID:AB_330985 |
| Rabbit monoclonal anti-phospho-p70 S6 Kinase (Thr389) (clone 108D2) | Cell Signaling Technology | Cat#9234<br>RRID:AB_2269803 |
| Rabbit monoclonal anti-Cyclin D1 (clone E3P5S) | Cell Signaling Technology | Cat#55506<br>RRID:AB_2827374 |
| Mouse monoclonal anti-Cyclin D3 (clone DCS22) | Cell Signaling Technology | Cat#2936<br>RRID:AB_2070801 |
| Rabbit monoclonal anti-phospho-Rb (Ser807/811) (clone D20B12) | Cell Signaling Technology | Cat#8516<br>RRID:AB_11178658 |
| Rabbit polyclonal anti-Cleaved Caspase-3 (Asp175) | Cell Signaling Technology | Cat#9661<br>RRID:AB_2341188 |
| Rabbit monoclonal anti-Cleaved PARP (Asp214) (clone D64E10) | Cell Signaling Technology | Cat#5625<br>RRID:AB_10699459 |
| Rabbit monoclonal anti-β-Actin (clone 13E5) | Cell Signaling Technology | Cat#4970<br>RRID:AB_2223172 |
| Rabbit polyclonal anti-Rb (Ser780) | Cell Signaling Technology | Cat#9307<br>RRID:AB_330015 |
| Mouse monoclonal anti-SYP (clone Snp-88) | BioGenex | Cat#MU363-UC<br>RRID:AB_2847824 |
| Mouse monoclonal anti-CDX2 (clone CDX2-88) | BioGenex | Cat#MU392A<br>RRID:AB_2923402 |
| **Biological samples** |  |  |
| AC patient-derived organoids | This study | Table S1 |
| CRC patient-derived organoids | Moorman et al. | N/A |
| **Chemicals, peptides, and recombinant proteins** |  |  |
| Advanced DMEM/F12 | Thermo Fisher Scientific | Cat#12634010 |
| GlutaMAX™ Supplement | Thermo Fisher Scientific | Cat#35050061 |
| HEPES | Thermo Fisher Scientific | Cat#15630106 |
| N-acetyl-L-cysteine | Sigma-Aldrich | Cat#A7250 |
| B27 supplement | Thermo Fisher Scientific | Cat#17504044 |
| Primocin® | InvioGen | Cat#ant-pm |
| Penicillin-Streptomycin | Thermo Fisher Scientific | Cat#15140148 |
| Amphotericin B | Cytiva | Cat# SV30078 |
| Nystatin | Millipore Sigma | Cat#N9150 |
| DNase I | Millipore Sigma | Cat#AMPD1 |
| Tumor Dissociation Kit | Milltenyi Biotec | Cat#130-095-929 |
| TrypLE™ | Thermo Fisher Scientific | Cat#12604013 |
| TheraPEAK™ ACK Lysing Buffer | Lonza | Cat#BP10-548E |
| Noggin-Fc Fusion Protein Conditioned Medium | IPA Therapeutics | Cat#N002 |
| Human IGF-I Recombinant Protein | PeproTech | Cat#100-11 |
| A83-01 | Millipore | Cat#SML0788 |
| Human EGF Recombinant Protein | PeproTech | Cat#100-15 |
| Human FGF2 Recombinant Protein | PeproTech | Cat#100-18B |
| Wnt Surrogate-Fc Fusion Protein | IPA Therapeutics | Cat#N001 |
| 5-FU | Selleckchem | Cat#S8848 |
| Oxaliplatin | MedChemExpress | Cat#HY-17371 |
| Mitomycin C | Fisher Scientific | Cat#32-581-0 |
| Futibatinib | Selleckchem | Cat#S8848 |
| TokeOni | Millipore Sigma | Cat#808350 |
| PEG-400 | Sigma-Aldrich | Cat#8.07485 |
| Solutol® | Sigma-Aldrich | Cat#42966 |
| Universal HIER antigen retrieval reagent (10X) | Abcam | Cat#ab208572 |
| **Critical commercial assays** |  |  |
| CellTiter-Glo 3D Cell Viability Assay | Promega | Cat#G9683 |
| PicoPure™ DNA Extraction kit | Thermo Fisher Scientific | Cat#KIT0103 |
| **Experimental models: Cell lines** |  |  |
| AC patient-derived organoids | This study | Table S1 |
| CRC patient-derived organoids | Moorman et al.<sup>43</sup> | N/A |
| **Experimental models: Organisms/strains** |  |  |
| Mouse: NOD.Cg-Prkdcscid Il2rgtm1Wjl/SzJ | The Jackson Laboratory | RRID:IMSR_JAX:005557 |
| **Recombinant DNA** |  |  |
| pLenti-PGK-tdTomato-AkaLuc | Moorman et al.<sup>43</sup> | N/A |
| **Software and algorithms** |  |  |
| Graphpad Prism 10 | GraphPad Software | https://www.graphpad.com/ |
| Cell Ranger 5.0.1 | 10X Genomics | https://support.10xgenomics.com/single-cell-gene-expression/software/pipelines/latest/what-is-cell-ranger |
| scVI tools 0.20.3 | Lopez et al.<sup>87</sup> | https://github.com/scverse/scvi-tools |
| QuPath-0.5.1-x64 | Bankhead et al.<sup>88</sup> | https://qupath.github.io/ |

## References

1. Welten, V. M. & Melnitchouk, N. Management of Peritoneal Disease in Colorectal Cancer. Hematol Oncol Clin North Am 36, 569–582 (2022).

2. Mei, S., Chen, X., Wang, K. & Chen, Y. Tumor microenvironment in ovarian cancer peritoneal metastasis. Cancer Cell International 2023 23:1 23, 1–13 (2023).

3. Gwee, Y. X. et al. Integration of Genomic Biology Into Therapeutic Strategies of Gastric Cancer Peritoneal Metastasis. Journal of Clinical Oncology 71, (2022).

4. Huang, Y., Alzahrani, N. A., Chua, T. C., Liauw, W. & Morris, D. L. Impacts of peritoneal cancer index on the survival outcomes of patients with colorectal peritoneal carcinomatosis. International Journal of Surgery 32, 65–70 (2016).

5. Verwaal, V. J. et al. Long-term survival of peritoneal carcinomatosis of colorectal origin. Ann Surg Oncol 12, 65–71 (2005).

6. Zhu, Y., Hanna, N., Boutros, C. & Alexander, H. R. Assessment of clinical benefit and quality of life in patients undergoing cytoreduction and Hyperthermic Intraperitoneal Chemotherapy (HIPEC) for management of peritoneal metastases. J Gastrointest Oncol 4, 62–71 (2013).

7. Franko, J. et al. Treatment of colorectal peritoneal carcinomatosis with systemic chemotherapy: A pooled analysis of North Central Cancer Treatment Group phase III trials N9741 and N9841. Journal of Clinical Oncology 30, 263–267 (2012).

8. Kemeny, N. E. et al. Updated long-term survival for patients with metastatic colorectal cancer treated with liver resection followed by hepatic arterial infusion and systemic chemotherapy. J Surg Oncol 113, 477–484 (2016).

9. D, B., S, K., B, L., MR, B. & M, D. Early and long-term postoperative management following cytoreductive surgery and hyperthermic intraperitoneal chemotherapy. World J Gastrointest Oncol 2, 36 (2010).

10. Mehta, S. S., Gelli, M., Agarwal, D. & Goéré, D. Complications of Cytoreductive Surgery and HIPEC in the Treatment of Peritoneal Metastases. Indian J Surg Oncol 7, 225 (2016).

11. Grotz, T. E. et al. Cytoreductive Surgery and Hyperthermic Intraperitoneal Chemotherapy for Moderately and Poorly Differentiated Appendiceal Adenocarcinoma: Survival Outcomes and Patient Selection. Ann Surg Oncol 24, 2646–2654 (2017).

12. Acs, M. et al. Prolonged Survival in Peritoneal Metastatic Appendiceal Carcinoma Patients Treated With Combined Cytoreductive Surgery and Hyperthermic Intraperitoneal Chemotherapy. Journal of Surgical Research 283, 839–852 (2023).

13. Osterlund, P. et al. Repeated centralized multidisciplinary team assessment of resectability, clinical behavior, and outcomes in 1086 Finnish metastatic colorectal cancer patients (RAXO): A nationwide prospective intervention study. The Lancet Regional Health - Europe 3, 100049 (2021).

14. Riihimäki, M., Hemminki, A., Sundquist, K., Sundquist, J. & Hemminki, K. Metastatic spread in patients with gastric cancer. Oncotarget 7, 52307 (2016).

15. Votanopoulos, K. I., Shen, P., Skardal, A. & Levine, E. A. Peritoneal Metastases from Appendiceal Cancer. Surg Oncol Clin N Am 27, 551 (2018).

16. Minhas, A., Hendrickson, J. & Minhas, S. A. Frequency and Risk Factors for Metastasis in Newly Diagnosed Appendiceal Carcinoma. Cureus 13, (2021).

17. Cortés-Guiral, D. et al. Primary and metastatic peritoneal surface malignancies. Nature Reviews Disease Primers 2021 7:*1* 7, 1–23 (2021).

18. Mikuła-Pietrasik, J., Uruski, P., Tykarski, A. & Książek, K. The peritoneal “soil” for a cancerous “seed”: a comprehensive review of the pathogenesis of intraperitoneal cancer metastases. Cellular and Molecular Life Sciences 2017 75:*3* 75, 509–525 (2017).

19. Tan, D. S., Agarwal, R. & Kaye, S. B. Mechanisms of transcoelomic metastasis in ovarian cancer. Lancet Oncology 7, 925–934 (2006).

20. Strach, M. C., Sutherland, S., Horvath, L. G. & Mahon, K. The role of chemotherapy in the treatment of advanced appendiceal cancers: summary of the literature and future directions. Ther Adv Med Oncol 14, 17588359221112478 (2022).

21. Dilly, A., Honick, B. D., Lee, Y. J., Bartlett, D. L. & Choudry, H. A. Rational application of targeted therapeutics in mucinous colon/appendix cancers with positive predictive factors. Cancer Med 9, 1753– 1767 (2020).

22. Shen, J. P. et al. Efficacy of Systemic Chemotherapy in Patients With Low-grade Mucinous Appendiceal Adenocarcinoma: A Randomized Crossover Trial. JAMA Netw Open 6, e2316161–e2316161 (2023).

23. Foote, M. B. et al. Molecular Classification of Appendiceal Adenocarcinoma. Journal of Clinical Oncology 41, 1553–1564 (2023).

24. Son, I. T. et al. Comparison of long-term oncological outcomes of appendiceal cancer and colon cancer: A multicenter retrospective study. Surg Oncol 25, 37–43 (2016).

25. Foote, M. B. et al. The Impact of Germline Alterations in Appendiceal Adenocarcinoma. Clinical Cancer Research 29, 2631–2637 (2023).

26. Aguirre, N. et al. Predictors of Recurrence in Nonmetastatic Appendiceal Epithelial Cancers: An Updated Single-Center Experience Over 25 Years. Ann Surg Oncol 32, 695–702 (2024).

27. Yaeger, R. et al. Clinical Sequencing Defines the Genomic Landscape of Metastatic Colorectal Cancer. Cancer Cell 33, 125–136.e3 (2018).

28. Afolabi, H. A. et al. A GNAS Gene Mutation’s Independent Expression in the Growth of Colorectal Cancer: A Systematic Review and Meta-Analysis. Cancers (Basel) 14, 5480 (2022).

29. Votanopoulos, K. I. et al. Appendiceal Cancer Patient-Specific Tumor Organoid Model for Predicting Chemotherapy Efficacy Prior to Initiation of Treatment: A Feasibility Study. Ann Surg Oncol 26, 139–147 (2019).

30. Weitz, J. et al. An Ex Vivo Organotypic Culture Platform for Functional Interrogation of Human Appendiceal Cancer Reveals a Prominent and Heterogenous Immunological Landscape. Clinical Cancer Research 28, 4793–4806 (2022).

31. Ito, I. et al. Development and characterization of orthotopic patient-derived xenograft models of peritoneal metastatic mucinous appendiceal adenocarcinoma. ESMO Gastrointestinal Oncology 7, 100133 (2025).

32. Carr, N. J. et al. The histopathological classification, diagnosis and differential diagnosis of mucinous appendiceal neoplasms, appendiceal adenocarcinomas and pseudomyxoma peritonei. Histopathology 71, 847–858 (2017).

33. Hoehn, R. S. et al. Current Management of Appendiceal Neoplasms. American Society of Clinical Oncology Educational Book 118–132 (2021) doi:10.1200/EDBK_321009.

34. Connor, S. J., Hanna, G. B. & Frizelle, F. A. Retrospective clinicopathologic analysis of appendiceal tumors from 7,970 appendectomies. Dis Colon Rectum 41, 75–80 (1998).

35. Reid, M. D. et al. Adenocarcinoma ex-goblet cell carcinoid (appendiceal-type crypt cell adenocarcinoma) is a morphologically distinct entity with highly aggressive behavior and frequent association with peritoneal/intra-abdominal dissemination: an analysis of 77 cases. Modern Pathology 2016 29:*10* 29, 1243–1253 (2016).

36. Boegh, M. & Nielsen, H. M. Mucus as a Barrier to Drug Delivery – Understanding and Mimicking the Barrier Properties. Basic Clin Pharmacol Toxicol 116, 179–186 (2015).

37. Kufe, D. W. Mucins in cancer: function, prognosis and therapy. Nature Reviews Cancer 2009 9:*12* 9, 874–885 (2009).

38. Kalra, A. V. & Campbell, R. B. Mucin impedes cytotoxic effect of 5-FU against growth of human pancreatic cancer cells: overcoming cellular barriers for therapeutic gain. British Journal of Cancer 2007 97:*7* 97, 910–918 (2007).

39. Boegh, M. & Nielsen, H. M. Mucus as a Barrier to Drug Delivery – Understanding and Mimicking the Barrier Properties. Basic Clin Pharmacol Toxicol 116, 179–186 (2015).

40. de Bruijn, I. et al. Analysis and Visualization of Longitudinal Genomic and Clinical Data from the AACR Project GENIE Biopharma Collaborative in cBioPortal. Cancer Res 83, 3861–3867 (2023).

41. Gao, J. et al. Integrative analysis of complex cancer genomics and clinical profiles using the cBioPortal. Sci Signal 6, (2013).

42. Cerami, E. et al. The cBio cancer genomics portal: an open platform for exploring multidimensional cancer genomics data. Cancer Discov 2, 401–404 (2012).

43. Moorman, A. R. et al. Progressive plasticity during colorectal cancer metastasis. Nature 2024 1–8 (2024) doi:10.1038/s41586-024-08150-0.

44. Hanahan, D. Hallmarks of Cancer: New Dimensions. Cancer Discov 12, 31–46 (2022).

45. Hanahan, D. & Weinberg, R. A. Hallmarks of cancer: The next generation. Cell 144, 646–674 (2011).

46. Hanahan, D. & Weinberg, R. A. The Hallmarks of Cancer. Cell 100, 57–70 (2000).

47. Fujii, M. et al. Human Intestinal Organoids Maintain Self-Renewal Capacity and Cellular Diversity in Niche-Inspired Culture Condition. Cell Stem Cell 23, 787–793.e6 (2018).

48. Ang, C. S.-P. et al. Genomic Landscape of Appendiceal Neoplasms. JCO Precis Oncol 2, PO.17.00302 (2018).

49. Bell, D. et al. Integrated genomic analyses of ovarian carcinoma. Nature 474, 609–615 (2011).

50. Sanchez-Vega, F. et al. Oncogenic Signaling Pathways in The Cancer Genome Atlas. Cell 173, 321–337.e10 (2018).

51. Fearon, E. R. & Vogelstein, B. A genetic model for colorectal tumorigenesis. Cell 61, 759–767 (1990).

52. Kim, G. P. et al. Phase III noninferiority trial comparing irinotecan with oxaliplatin, fluorouracil, and leucovorin in patients with advanced colorectal carcinoma previously treated with fluorouracil: N9841. J Clin Oncol 27, 2848–2854 (2009).

53. Hurwitz, H. et al. Bevacizumab plus irinotecan, fluorouracil, and leucovorin for metastatic colorectal cancer. N Engl J Med 350, 2335–2342 (2004).

54. Goldberg, R. M. et al. A randomized controlled trial of fluorouracil plus leucovorin, irinotecan, and oxaliplatin combinations in patients with previously untreated metastatic colorectal cancer. J Clin Oncol 22, 23–30 (2004).

55. Franko, J. et al. Treatment of colorectal peritoneal carcinomatosis with systemic chemotherapy: A pooled analysis of North Central Cancer Treatment Group phase III trials N9741 and N9841. Journal of Clinical Oncology 30, 263–267 (2012).

56. Holderfield, M. et al. Concurrent inhibition of oncogenic and wild-type RAS-GTP for cancer therapy. Nature 629, 919–926 (2024).

57. Jiang, J. et al. Translational and Therapeutic Evaluation of RAS-GTP Inhibition by RMC-6236 in RAS-Driven Cancers. Cancer Discov 14, 994–1017 (2024).

58. Rialdi, A. et al. WNTinib is a multi-kinase inhibitor with specificity against β-catenin mutant hepatocellular carcinoma. Nature Cancer 2023 4:*8* 4, 1157–1175 (2023).

59. Fujii, M. et al. Human Intestinal Organoids Maintain Self-Renewal Capacity and Cellular Diversity in Niche-Inspired Culture Condition. Cell Stem Cell 23, 787–793.e6 (2018).

60. Zhao, Q. et al. FGFR inhibitor, AZD4547, impedes the stemness of mammary epithelial cells in the premalignant tissues of MMTV-ErbB2 transgenic mice. Sci Rep 7, (2017).

61. Meric-Bernstam, F. et al. Futibatinib, an Irreversible FGFR1-4 Inhibitor, in Patients with Advanced Solid Tumors Harboring FGF/ FGFR Aberrations: A Phase I Dose-Expansion Study. Cancer Discov 12, 402– 415 (2022).

62. Pretzsch, E. et al. Mechanisms of Metastasis in Colorectal Cancer and Metastatic Organotropism: Hematogenous versus Peritoneal Spread. J Oncol 2019, 7407190 (2019).

63. Kanda, M. & Kodera, Y. Molecular mechanisms of peritoneal dissemination in gastric cancer. http://www.wjgnet.com/ 22, 6829–6840 (2016).

64. Mikuła-Pietrasik, J., Uruski, P., Tykarski, A. & Książek, K. The peritoneal “soil” for a cancerous “seed”: a comprehensive review of the pathogenesis of intraperitoneal cancer metastases. Cellular and Molecular Life Sciences 2017 75:*3* 75, 509–525 (2017).

65. Ganesh, K. et al. A rectal cancer organoid platform to study individual responses to chemoradiation. Nature Medicine 2019 25:*10* 25, 1607–1614 (2019).

66. Van De Wetering, M. et al. Prospective derivation of a Living Organoid Biobank of colorectal cancer patients. Cell 161, 933 (2015).

67. Vlachogiannis, G. et al. Patient-derived organoids model treatment response of metastatic gastrointestinal cancers. Science 359, 920 (2018).

68. Sachs, N. et al. A Living Biobank of Breast Cancer Organoids Captures Disease Heterogeneity. Cell 172, 373–386.e10 (2018).

69. Kitai, H. et al. Combined inhibition of KRASG12C and mTORC1 kinase is synergistic in non-small cell lung cancer. Nature Communications 2024 15:*1* 15, 1–15 (2024).

70. She, Q. B. et al. 4E-BP1 Is a Key Effector of the Oncogenic Activation of the AKT and ERK Signaling Pathways that Integrates Their Function in Tumors. Cancer Cell 18, 39–51 (2010).

71. Yao, H. & He, S. Multi-faceted role of cancer-associated adipocytes in the tumor microenvironment. Mol Med Rep 24, 866 (2021).

72. Quail, D. F. & Dannenberg, A. J. The obese adipose tissue microenvironment in cancer development and progression. Nat Rev Endocrinol 15, 139–154 (2019).

73. Gil, A., Olza, J., Gil-Campos, M., Gomez-Llorente, C. & Aguilera, C. M. Is adipose tissue metabolically different at different sites? International Journal of Pediatric Obesity 6, 13–20 (2011).

74. Rijken, A. et al. The Burden of Peritoneal Metastases from Gastric Cancer: A Systematic Review on the Incidence, Risk Factors and Survival. J Clin Med 10, 4882 (2021).

75. Roth, L. et al. Peritoneal Metastasis: Current Status and Treatment Options. Cancers (Basel) 14, 60 (2021).

76. Huang, Y., Alzahrani, N. A., Chua, T. C., Liauw, W. & Morris, D. L. Impacts of peritoneal cancer index on the survival outcomes of patients with colorectal peritoneal carcinomatosis. International Journal of Surgery 32, 65–70 (2016).

77. Odendahl, K. et al. Quality of life of patients with end-stage peritoneal metastasis treated with Pressurized IntraPeritoneal Aerosol Chemotherapy (PIPAC). European Journal of Surgical Oncology (EJSO) 41, 1379–1385 (2015).

78. Verwaal, V. J. et al. Long-term survival of peritoneal carcinomatosis of colorectal origin. Ann Surg Oncol 12, 65–71 (2005).

79. Torre, L. A. et al. Ovarian cancer statistics, 2018. CA Cancer J Clin 68, 284–296 (2018).

80. Vaughan, S. et al. Rethinking Ovarian Cancer: Recommendations for Improving Outcomes. Nat Rev Cancer 11, 719 (2011).

81. Ganesh, K. et al. L1CAM defines the regenerative origin of metastasis-initiating cells in colorectal cancer. Nat Cancer 1, 28–45 (2020).

82. Cheng, D. T. et al. Memorial sloan kettering-integrated mutation profiling of actionable cancer targets (MSK-IMPACT): A hybridization capture-based next-generation sequencing clinical assay for solid tumor molecular oncology. Journal of Molecular Diagnostics 17, 251–264 (2015).

83. Chakravarty, D. et al. OncoKB: A Precision Oncology Knowledge Base. JCO Precis Oncol 1–16 (2017) doi:10.1200/PO.17.00011/SUPPL_FILE/DS_17.00011.DOCX.

84. OncoKB^TM^ Cancer Gene List. https://www.oncokb.org/cancer-genes.

85. Luecken, M. D. et al. Benchmarking atlas-level data integration in single-cell genomics. Nature Methods 2021 19:*1* 19, 41–50 (2021).

86. Xu, C. et al. Probabilistic harmonization and annotation of single-cell transcriptomics data with deep generative models. Mol Syst Biol 17, (2021).

87. Lopez, R., Regier, J., Cole, M. B., Jordan, M. I. & Yosef, N. Deep generative modeling for single-cell transcriptomics. Nature Methods 2018 15:*12* 15, 1053–1058 (2018).

88. Bankhead, P. et al. QuPath: Open source software for digital pathology image analysis. Scientific Reports 2017 7:*1* 7, 1–7 (2017).

